# Thalamo-hippocampal pathway regulates incidental memory load in mice

**DOI:** 10.1101/2021.08.07.453742

**Authors:** G. Torromino, V. Loffredo, D. Cavezza, G. Sonsini, F. Esposito, A. H. Crevenna, M. Gioffrè, M. De Risi, A. Treves, M. Griguoli, E. De Leonibus

## Abstract

Incidental memory can be challenged by increasing either the retention delay or the memory load. The dorsal hippocampus (dHP) appears to help with both consolidation from short-term (STM) to long-term memory (LTM), and higher memory loads, but the mechanism is not fully understood. Here we find that female mice, despite having the same STM capacity of 6 objects and higher resistance to distraction in our different object recognition task (DOT), when tested over 1h or 24h delays appear to transfer to LTM only 4 objects, whereas male mice have an STM capacity of 6 objects in this task. In male mice the dHP shows greater activation (as measured by c-Fos expression), whereas female mice show greater activation of the ventral midline thalamus (VMT). Optogenetic inhibition of the VMT-dHP pathway during off-line memory consolidation enables 6-object LTM retention in females, while chemogenetic VMT-activation impairs it in males. Thus, removing or enhancing sub-cortical inhibitory control over the hippocampus, leads to differences in incidental memory.

**One Sentence Summary:** The sex-dependent recruitment of a subcortical pathway regulates how many items are spontaneously memorized in mice.

## Introduction

*Memory* can be challenged by lengthening the retention interval (its *duration*) but also by augmenting what has to be retained (its *load*). Additional neural mechanisms are thought to be necessary as the required *duration* extends from short (seconds/minutes) to long (hours/days) and to remote (months/years) time scales, based on the evidence that the passage of time promotes the reorganization of neuronal circuits and activates the molecular processes referred to as memory consolidation. The dorsal hippocampus (dHP) has been involved in memory consolidation, the stabilization of a trace from short-term (STM) to long-term memory (LTM)^1^, and in spatial novelty detection^2,3^.

However, emerging evidence suggests that also heavier loads, quantified by the number of items that are to be stored in memory, lead to a reorganization of neuronal circuits and to distinct molecular processes. Converging findings in rodents, primates and humans suggest that high memory load drives dHP recruitment in STM item tasks. We and others have previously reported that, in male mice, object STM capacity appears limited to about 6 elements^4–7^. These findings have been obtained by using the Different/Identical Object Task (DOT/IOT), a modified version of the spontaneous object recognition task, which stimulates incidental encoding based on the spontaneous interest of animals to explore novel objects. In the DOT/IOT, the memory load is manipulated by increasing the number of different objects to remember during the study phase (e.g., 3, 4, 6 and 9 different objects). Using the DOT/IOT, we showed that male mice can perform the task in the short-term delay (1 min) for 3, 4 and 6 DOT, but not for 9; on the contrary, in the control task with identical objects (IOT) males can recognize the new object compared to all the familiar ones. Additionally, we also showed that dorsal hippocampal (dHP) lesions selectively impaired performance in the 6-DOT, but not the 4-DOT. These findings are in line with those obtained with the same or different tasks in male mice, rats or monkeys showing that lesion of the dHP reduces item memory capacity^4–6,8,9^. Interestingly, they are also in line with studies in humans showing that patients with hippocampal lesions show reduced memory span for items^10,11^.

How the number of elements to remember influences the formation of LTM, namely how duration and load interact, remains a relevant unexplored issue. This is likely because LTM is thought to have almost unlimited capacity, thanks to the possibility of prolonged or multiple trials of learning that facilitate consolidation. However, most of our daily life events are unique experiences, leading to incidental encoding, without necessarily the repetition needed to form stable memories.

Therefore, how the number of elements encountered during incidental encoding affects memory consolidation is an open question. Interestingly, in male mice we have reported that all the 6 objects encoded at short delay were also remembered at longer delays, through the activation of protein kinase A and Ca^2+^/calmodulin-dependent protein kinase (PKA/CaMKII) mediated GluR1 AMPA receptors phosphorylation at the serine sites S831 and S845 and *de novo* protein synthesis^8^. These findings in males suggested that all objects encoded in STM are also automatically consolidated into LTM.

While studying memory capacity in female mice by subjecting them to the DOT/IOT, however, we discovered that, as males they remember up to 6 different objects, but not 9 at short delay (1 min). However, differently from males, they spontaneously remember at long-delays (1 or 24 h) only 4 rather than the 6 objects encoded at short delay (1 min), although they are more resistant than males to retroactive interference at short delay (1 min). Though these sex differences do not probably reflect any major qualitative difference in neural mechanism, as they might originate from the use of different cognitive strategies to solve the task. However, they offer a precious window to shed light on the dynamic activation of neural circuits, and to study how the duration of the task, far from simply extending the retention of the incidentally acquired items, effectively restricts memory capacity. How does that come about?

Sex differences in cognitive functions is an emerging field. In the memory domain, sex differences have been documented^12–14^, and multiple lines of evidence suggest that in adulthood females tend to engage the dHP less than males, while they use more non-hippocampal dependent navigation strategies when they are engaged in spatial tasks. Here, we have tested, in mice, the hypothesis that sex differences in the dynamic recruitment of the dHP would influence high load incidental LTM. Additionally, we have used the observed sex differences in incidental LTM to provide mechanistic evidence on what drives dHP activation in conditions of high memory load, to help clarify this understudied question in the neurobiology of memory.

## Results

### Females transfer in LTM only 4 of the 6 objects encoded in STM

To set up the DOT/IOT in female mice, we have exposed them to the 3-, 4-, 6- and 9-different objects (DOT) or identical objects (IOT), using the same procedure previously used for male mice^4,8,19^ (Fig.1). We found that also females, like males, could remember up to six different objects, but not nine, at short-retention intervals (1 min) (Fig.1 a-d and Supplementary Table 1 and 5 for statistical analysis). However, when female mice were tested 1 or 24 h after the study phase, their incidental LTM memory span was limited to 4 different objects (Fig.1e-h), unlike that of males, which was 6. This result seems to be independent on the phases of their estrous cycle phases (see methods and Supplementary Fig. 1a-c). Thus, while males spontaneously “transfer” all the objects encountered from STM to LTM (1 or 24 h), females do the same in condition of low (6-IOT) or intermediate (4-DOT) memory load, but apparently not in the highest memory load conditions (Fig. 2a). Note, though, that they explore the objects more than males during the study phase (Supplementary Table 2). It must be noted, however, that the exploration time in the study phase does not correlate with the percentage of new object exploration at the test phase neither in males [simple regression between T2 and % New in the 6-DOT at 1 min delay: p = 0.7104; at 24 h delay: p = 0.4775), nor in females [simple regression between T2 and % New in the 6-DOT at 1 min delay: p = 0.5478; at 24 h delay: p = 0.0705]. Therefore, in female mice memory stabilization seems not to be automatic but optional, suggesting the possibility of different strategies, or cognitive attitudes, reflected in the high load incidental LTM.

**Figure 1.**
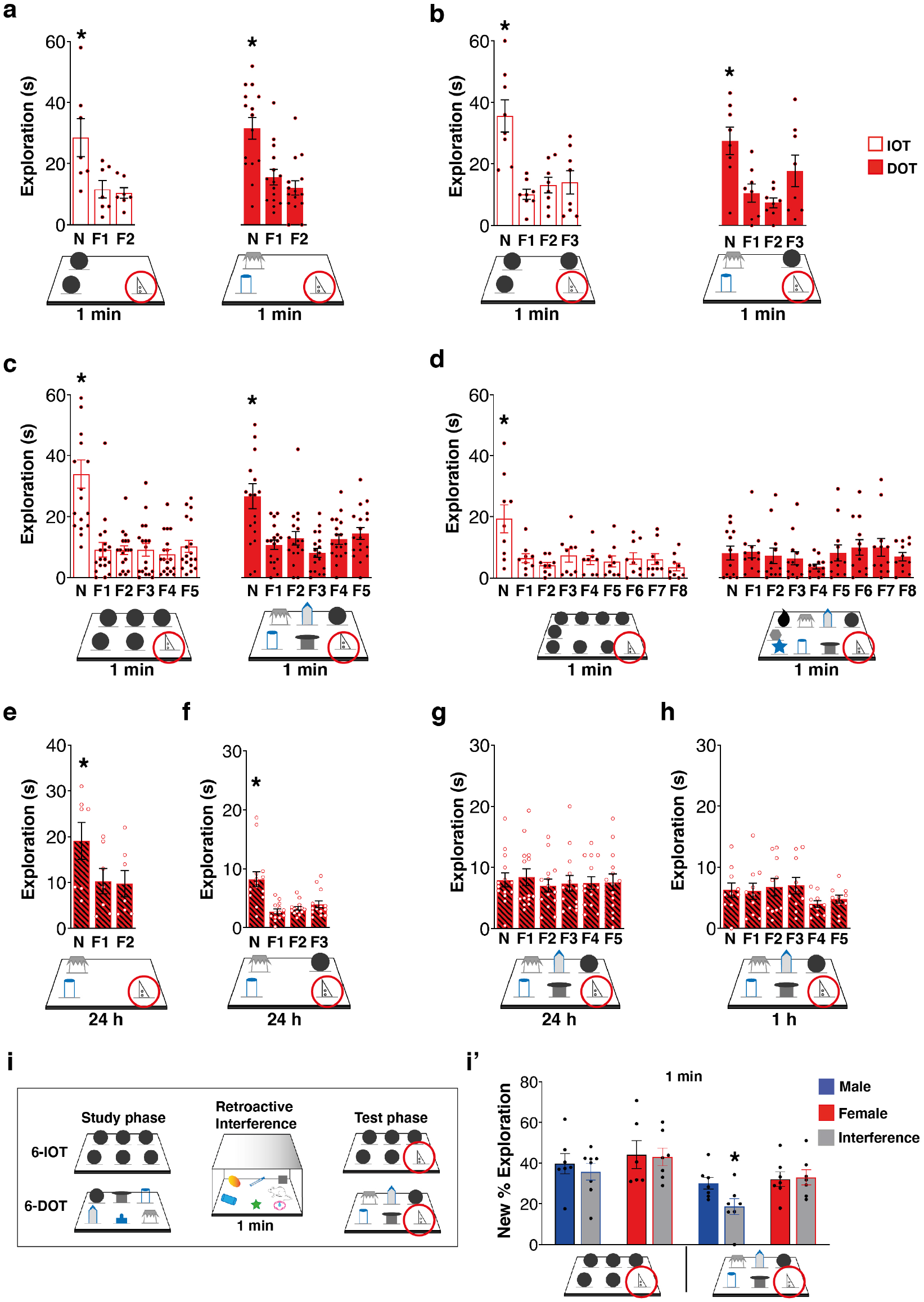
Females show a similar STM capacity to males, but greater resistance to retroactive interference at the expense of reduced memory consolidation. **(a-d)** Bar charts report exploration of the new object (circled) and each of the familiar ones (F) in the Different/Identical Object Task (DOT-IOT). Female mice explored significantly more the new object compared to all the familiar ones at 1 min delay in all the control tasks (IOTs) and in the 3-DOT (a, low load), 4-DOT (b, intermediate load) and 6-DOT (c, high load), but not in the 9-DOT (d, overload). **(e-h)** In the LTM tasks, females explored significantly more the new object compared to all the familiar ones at 24 h delay in the 3-DOT (e) and 4-DOT (f), but they showed an impaired performance in the 6-DOT (g), already at 1 h delay (h). * p < 0.05 New vs all the familiar objects (one-way repeated measure ANOVA followed by Dunnett post-hoc). **(i-i’)** Schematic of the task to test the effect of retroactive interference on memory capacity at 1 min delay (i). No effect of the interference protocol was found on the performance of males and females in the 6-IOT (i’). However, while interference caused a significant decrease of the new % exploration in males, no effect was revealed in females (i’), which demonstrated to be resistant to retroactive interference even in high memory load condition (6-DOT). * p < 0.05 control *vs* interference within sex (one-way ANOVA). Data in bar charts are presented as mean values ± SEM. For statistics see Supplementary Table 5.

**Figure 2.**
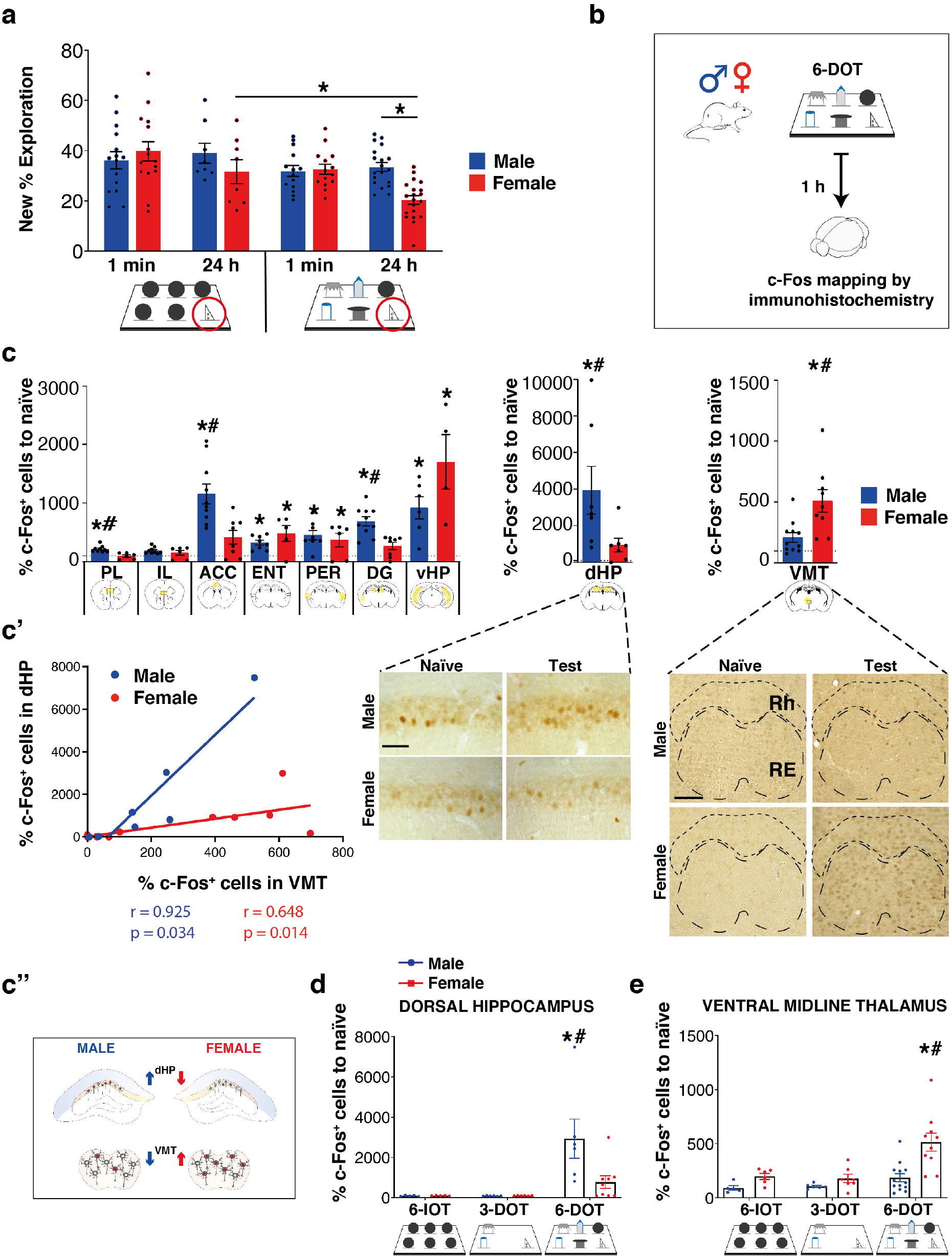
Females show an apparent capacity limit after memory consolidation, associated with VMT hyperactivation and dHP hypoactivation compared to males. **(a)** Bar charts report new object % exploration in the 6-IOT/DOT for STM (1 min) and LTM (24 h) of male (6-IOT: 1 min n = 15, 24 h n = 8; 6-DOT: 1 min n = 14, 24 h n = 19) and female (6-IOT: 1 min n = 15, 24 h n = 8; 6-DOT: 1 min n = 14, 24 h n = 20) mice. The performance of females in the 6-DOT at 24 h delay was significantly lower than that in the 6-IOT and that of males in the 6-DOT. **(b)** Schematic of the c-Fos mapping experiment. **(c)** Bar charts represent the mean of c-Fos^+^ cells normalized on naïve *per* sex for each brain region analyzed. The VMT was the brain region found significantly activated only in females exposed to the 6-DOT. The CA1-CA3 of the dHP was found significantly activated only in male mice exposed to the 6-DOT. The bottom panels show representative photographs of the dHP and the VMT in each experimental condition. Scale bar 50 µm. **(c’)** The Spearman correlation coefficient between VMT and dHP c-Fos^+^ cells was positive and significant for both sexes. **(c’’)** Illustration of the opposite pattern of activation between the dHP and the VMT in the two sexes. **(d-e)** Load-dependent activation of c-Fos in the VMT and the dHP for both sexes. The VMT was significantly more active in female mice after the 6-DOT compared to lower load conditions (6-IOT and 3-DOT) and to males (d). The same happens to the dHP of males (e). Data are normalized on naïve c-Fos expression. * p < 0.05 naïve vs test (within sexes), # p < 0.05 test between sexes (Bonferroni correction). Data in bar charts are presented as mean values ± SEM. For statistics see Supplementary Table 5 and Supplementary Fig. 2. PL = prelimbic cortex; IL = infralimbic cortex; ACC = anterior cingulate cortex; VMT = ventral midline thalamus (including the reuniens and the rhomboid thalamic nuclei); ENT = entorhinal cortex; PER = perirhinal cortex; dHP = dorsal hippocampus; DG = dentate gyrus; vHP = ventral hippocampus.

The engagement of LTM mechanisms in condition of high load has been shown to be more resistant than STM to retroactive interference^20^. Therefore, we tested “resistance to interference” in both sexes by introducing into the waiting cage a few distracting stimuli (such as other objects and biologically relevant odors) during the 1 min delay (Fig.1i), a procedure hereon called interference. Exposure to interference disrupted performance in males only in the high load condition (6-DOT) while, surprisingly, it did not affect it at all in females (Fig. 1i’). These data suggest that females, during spontaneous encoding of many objects, can also activate stabilization mechanisms, but in a way that it can make their STM *more* resistant to retroactive interference and *less* resistant to the effects of time passing, compared to males. Since these sex differences do not probably reflect structural difference in memory capacity, we have used them as a model to shed light on the dynamic activation of neural circuits that support high load incidental LTM and how it is modulated by sex.

### Sex- and load-dependent activation of neuronal circuits

To determine how the number of objects presented during the study phase regulates c-Fos activation in a sex-dependent manner we perfomed c-Fos brain activation experiments. For this reason, animals were subjected to the familiarization session (empty arena) and the study sessions (6 different objects) before being sacrificed for the immunohistological analyses to avoid any possible interference of the test phase on c-Fos expression, where the two groups differ. To control for interference due to different exploration in male and female mice, we matched female *vs* males with similar exploration (as schematized in Fig. 2b) during the study phase (males: 102.2 s ± 8.0; females 89.6 s ± 3.9; one-way ANOVA for exploration time F_1,17_ = 1.575, p = 0.2264). When analyzing c-Fos^+^ cells, we found no differences in the basal level of activation between sexes in naïve mice (Supplementary Table 3). The c-Fos counting values of mice exposed to the study phase of the 6-different objects was normalized to that of naïve mice. c-Fos expression, as compared to naïve animals, was increased in all the brain regions analyzed except for the infralimbic cortex (IL, Fig. 2c and Supplementary Fig. 2b). The entorhinal (ENT, Fig. 2c and Supplementary Fig. 2d) and perirhinal (PER, Fig. 2c and Supplementary Fig. 2e) cortices, and the ventral hippocampus (vHP, Fig. 2c and Supplementary Fig. 2g) were similarly activated in both sexes. The PER and ENT cortices are already known to be pivotal for object recognition in low memory load conditions^21–23^. Consistently, they were similarly activated for the two sexes, which makes highly unlikely their specific involvement in the observed sex differences in memory consolidation in high memory load conditions.

The 6-different objects induced a significantly higher activation in males than in females for the dHP (Fig. 2c), the dentate gyrus (DG, Fig. 2c and Supplementary Fig. 2f), the prelimbic (PL, Fig. 2c and Supplementary Fig. 2a) and anterior cingulate cortices (ACC, Fig. 2c and Supplementary Fig. 2c). On the contrary, the 6-different objects induced higher activation in females than in males only in the ventral thalamic nuclei (VMT), including the reuniens (RE) and the rhomboid nucleus (RH) (VMT, Fig. 2c), which have similar anatomical connections with the HP and exert similar functional modulation of cortical-to-cortical communication required for system consolidation^1^.

We focused on brain regions that showed opposite patterns of activation in males and females after the 6-different objects, specifically on the dHP and the VMT, and not on the ACC, as chemogenetic inhibition of the ACC using *Designer Receptors Exclusively Activated by Designer Drugs* (DREADDs) during early memory consolidation, proved not to affect 6-DOT memory performance at 24 h delay (Supplementary Fig. 3).

**Figure 3.**
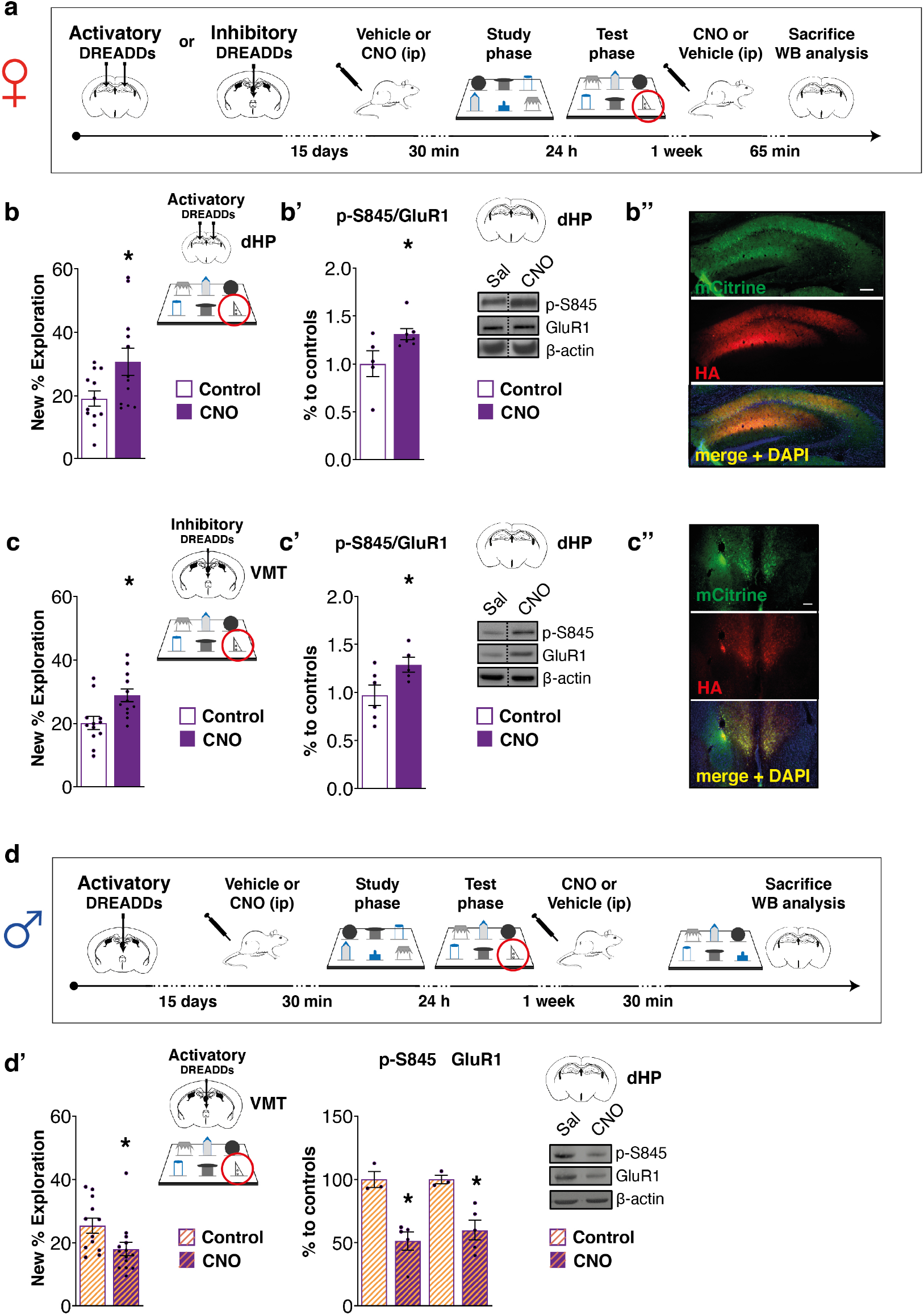
Activation of the dHP and inhibition of the VMT rescues the expression of high load incidental LTM. **(a)** Schematic of the experimental design showing the timing of injection of an activatory DREADDs in the dHP or an inhibitory one in the VMT of female mice and the experimental course. **(b)** AAV-CaMKIIa-HA-rM3D(Gs)-IRES-mCitrine was injected in the dHP of females. Bar chart represents increased new object % exploration for CNO treated females compared to controls. **(b’)** Bar charts represent western blot analysis of p-S845/GluR1 levels in the dHP of vehicle and CNO treated mice, which is increased by dHP chemogenetic activation. No change was detected in the expression of GluR1. On the right are shown representative bands for each condition. **(b’’)** Representative images of AAV expression in the dHP; scale bar 100 µm. **(c)** AAV-CaMKIIa-HA-rM4D(Gi)-IRES-mCitrine was injected in the VMT of female mice. Bar charts represent increased new object % exploration in CNO treated females compared to controls. **(c’)** Bar charts represent western blot analysis of p-S845/GluR1 levels in the hippocampus of vehicle and CNO treated mice, which shows a significant increase after VMT inhibition. Concomitant we found that VMT inhibition also increased the GluR1 levels. On the right are shown representative bands for each condition. **(c’’)** Representative images of AAV expression in the VMT; scale bar 100 µm.**(d)** Schematic of the experimental design showing the timing of injection of an activatory DREADDs in the VMT of male mice and the experimental course. **(d’)** AAV-CaMKIIa-HA-rM3D(Gs)-IRES-mCitrine was injected in the VMT of males. Bar charts on the left represent new object % exploration for vehicle and CNO treated mice showing that VMT activation induced a memory consolidation capacity limit in males. Bar charts on the right represent western blot analysis of p-S845 and GluR1 levels in the dHP of vehicle and CNO treated mice, which were both decreased by the chemogenetic activation of the VMT, while no change was detected on their ratio. Representative bands for each condition are shown on the right. * p < 0.05 between groups. Data in bar charts are presented as mean values ± SEM. Dashed lines in (b’) and (c’) delineate cut parts from the same western blot. For statistics see Supplementary Table 5.

To better correlate the object memory load with c-Fos activation we performed an additional experiment by evaluating the VMT and dHP c-Fos expression after exposing the animals to low (6-identical objects), intermediate (3-different objects) and high (6-different objects) memory load (Fig. 2d-e). Based on the existence of reciprocal connections between the dHP and the VMT (confirmed by us, as reported in Supplementary Fig. 4)^24,25^, we performed a Spearman correlational analysis of VMT and dHP activation after the 6-different objects, which resulted positive in both sexes (Fig. 2c’). The VMT-dHP correlation plots of males and females clearly showed that, although their activation was correlated in both cases, the average values were different between the two sexes: higher average VMT activation levels in females than males corresponded with lower activation values in the dHP (Fig. 2c’’). We found no correlation between the time spent exploring objects in the study phase and the c-Fos expression levels in the VMT or the dHP of males and females [simple regression between T2 and c-Fos in VMT: for males p = 0.5709, for females p = 0.2506; simple regression between T2 and c-Fos in dHP: for males p = 0.9616, for females p = 0.9868]. This is in line with our previous findings, namely that in males, object exploration during the study phase showed no correlation with discrimination during the test phase and / or GluA1 AMPA receptor phosphorylation in the dHP^8^. This finding supports the hypothesis that the different level of c-Fos activation in the two regions of interest in males and females was induced by “object coding in memory” rather than “object exploration”.

**Figure 4.**
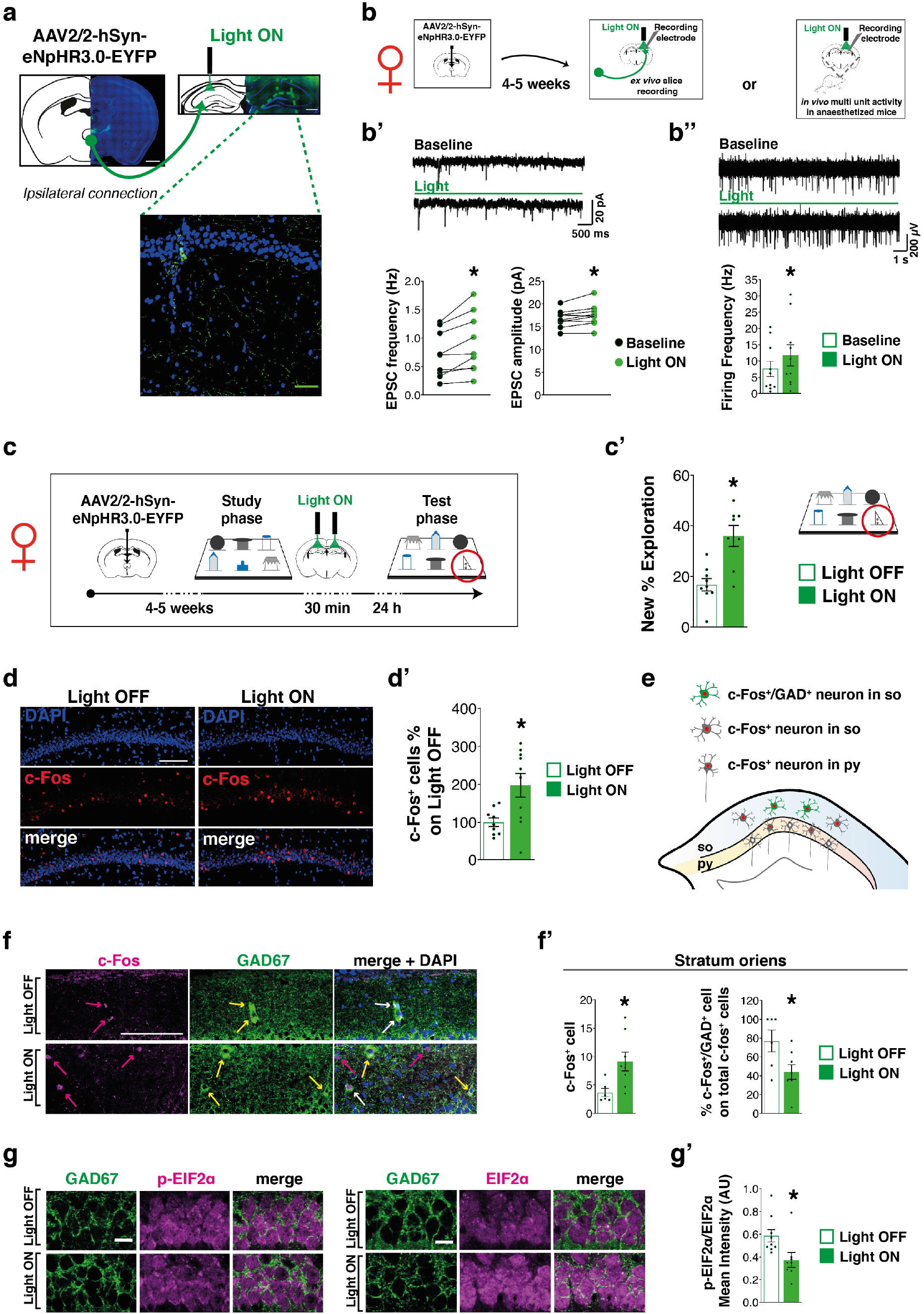
VMT-dHP photoinhibition increases hippocampal activity and rescues high load incidental LTM in females. **(a)** Representative images of the AAV injection in the VMT and its diffusion in the fiber terminals at the level of the dHP (scale bar 100 µm and 50 µm in the squared magnification). **(b)** Schematic of the experimental plans: AAV-hSyn-eNpHR3.0-EYFP was injected in the VMT and after about 4-5 weeks mice underwent *ex vivo* recordings in hippocampal slices (b’) or *in vivo* hippocampal multiunit activity recordings under anesthesia (b’’). **(b’)** On the top, representative traces from CA1 hippocampal neurons showing EPSCs in baseline and under photoinhibition (green bar). On the bottom, scatter plot showing EPSPs increased frequency and amplitude for females. **(b’’)** On the top, representative traces of multiunit activity recorded from hippocampal CA1 pyramidal layer before and during photoinhibition (green bar). On the bottom, bar charts represent increased frequency activity in females between baseline (control) and under photoinhibition (light ON). * p < 0.05 between groups (Wilcoxon test). **(c)** Schematic of the experimental design showing the timing of VMT-dHP fibers post-training (30 min) continuous inhibition. **(c’)** Bar charts represent the new object % exploration in light OFF and light ON females in the test conducted 24 h after post-training VMT-dHP photoinhibition, which shows a rescued memory consolidation capacity. **(d-d’)** VMT-dHP photoinhibition increased c-Fos^+^ cells in the CA1 pyramidal layer of females; scale bars: 100 µm. **(e)** Schematic of the dHP pyramidal layer (py) and *stratum oriens* (so) in which the c-Fos^+^ cells and the co-localizing signal with GAD67 (GAD^+^ cells) was analysed. **(f-f’)** VMT-dHP photoinhibition also increased c-Fos^+^ cells in the so of the CA1, but the percentage of GAD^+^/c-Fos^+^ cells was lowered by VMT-dHP photoinhibition; scale bar 100 µm; magenta arrows indicate c-Fos^+^ cells, yellow arrows GAD^+^ ones, while white arrows indicate c-Fos^+^/GAD^+^ cells. **(g-g’)** Quantitative analyses of immunofluorescence intensity in the CA1 excitatory neurons after VMT-dHP photoinhibition showed decreased p-EIF2α/EIF2α ratio in females; scale bar 10 µm. * p < 0.05 between groups. Data in bar charts are presented as mean values ± SEM. For statistics see Supplementary Table 5.

### Chemogenetic modulation of VMT removes the LTM limit

To provide a causal link between the observed activation of the VMT and dHP in regulating sex differences in memory capacity we chemogenetically and optogenetically manipulated these two brain areas.

It is known that the VMT can modulate hippocampal excitability^25–27^; thus we tested the hypothesis that VMT activation regulates the engagement/disengagement of the dHP in the spontaneous expression of memory observed with the DOT. Therefore, we first verified whether the hypoactivation of the dHP and hyperactivation of the VMT have a functional role in the DOT memory observed in females (Fig. 3a). The chemogenetic stimulation of the dHP (spearing the vHP, Supplementary Fig. 5) and inhibition of the VMT both removed the apparent capacity limit in females (Fig. 3b-b’’ and Fig. 3c-c’’) and, remarkably, increased the phosphorylation of the serine site S845 (p-S845) of the GluR1 subunit of the AMPA receptor in the dHP (Fig. 3b’-c’), which is a mechanism required for processing high memory load in males^8^. Importantly, this effect was the first evidence we obtained that the inhibition of the VMT could rescue the dHP activation in females (Fig. 3c’). We excluded that these effects were due to the injection of the clozapine-N-oxide (CNO) *per se*, as the same dose of CNO injected in naïve female mice did not affect either memory or p-S845 (Supplementary Fig. 6a-a’). Moreover, we showed that the dHP chemogenetic activation effect was time-dependent because injection of CNO immediately post-training was ineffective (Supplementary Fig. 6b-b’). This finding shows that in order to remove the apparent capacity limit, hippocampal activation must be induced in the early phase of memory consolidation. Considering the pharmacokinetic of CNO^28^, our experimental design was aimed at manipulating brain activation during off-line early memory consolidation, mostly not affecting performance during the study and test phases. Accordingly, when comparing the total exploration of males inoculated with the activatory DREADDs in the VMT and females inoculated with the inhibitory DREADDs in the VMT at the study phase of the 6-DOT, we found an effect for sex but not for treatment or interaction sex *vs* treatment [two way ANOVA: effect for sex F_1,46_ = 21.933, p < 0.0001; effect for treatment: F_1,46_ = 0.205, p = 0.6525; effect for sex *vs* treatment F_1,46_ = 0.006, p = 0.9411]. Of note, no changes were observed on the general level of exploration during the test phase (T3, Supplementary Table 4), suggesting that VMT manipulations were not changing the general level of exploration of the familiar objects, but specifically modulating the preference for the novel object during testing.

This comparison confirms our initial observation that females are more explorative than males in baseline conditions (see Supplementary Table 2) and it suggests that our chemogenetic manipulation of the VMT did not interfere with the general arousal level or exploratory behavior of mice, but acted mostly in the post-study phase by inhibiting or activating the VMT in response to the 6-DOT, during early off-line memory consolidation.

We then asked whether exogenous inhibition of the VMT could expand the memory consolidation capacity in male mice; to test this hypothesis we chemogenetically inhibited the VMT of a new group of males after exposition to an overload condition (8-DOT) and tested the animals 24 h later (Supplementary Fig. 7). Our results showed that this approach did not result in novel object recognition in the 8-DOT (Supplementary Fig. 7), and so that manipulation of the VMT-dHP does not expand memory beyond its normal limit.

### Inhibition of the VMT-dHP pathway removes the LTM limit

We next asked whether the limit on the spontaneous memory expression associated with the hyperactivation of the VMT is sex specific. To address this issue, we “feminized” the VMT of a group of male mice by chemogenetical stimulation (Fig. 3d). This approach led to the same apparent limit as that of females and decreased the level of p-S845 and GluR1 after exposure to the 6-DOT compared to controls (Fig. 3d’), without affecting the general exploratory activity of males (Supplementary Table 4). Similarly to previous results (see Supplementary Table 2), males were shown to be less explorative than females independently of the treatment received [two way ANOVA for total exploration at the study phase between males inoculate with an activatory DREADDs in the VMT and females inoculated with an inhibitory DREADDs in the VMT: effect for sex F_1,46_ = 21.933, p < 0.0001], suggesting that DREADDs were not modulating the general level of exploratory behavior.

The data obtained so far suggested that VMT hyperactivation negatively modulates the dHP, and that in females the high load of the 6-DOT naturally triggers this condition resulting in dHP hypoactivation. To confirm this hypothesis, we used an optogenetics approach aimed at selectively inhibiting the VMT-dHP direct pathway. We first injected an adeno-associated viral vector expressing Halorhodopsin (eNpHR3.0) into the VMT and showed that in females, continuous photoinhibition of the VMT post-learning was sufficient to rescue the expression of spontaneous memory in the 6-DOT (Supplementary Fig. 8a-a’). Then, we injected eNpHR3.0 in the VMT and placed the fiber optic cannulae into the dHP in order to inhibit specifically the VMT-dHP pathway (Fig. 4a, Supplementary Fig. 9a).

Using three parallel approaches, we showed that in females, photoinhibition of the VMT-dHP pathway: 1) increased the frequency and amplitude of the spontaneous excitatory post synaptic currents (EPSCs) in the dHP, as measured by *ex-vivo* patch clamp recordings in hippocampal slices (Fig. 4b-b’); 2) increased the spontaneous extracellular multiunit activity (MUA) in the pyramidal layer of the CA1 region recorded *in vivo* in anaesthetized mice (Fig. 4b-b’’); 3) removed the apparent memory limit in the 6-DOT (Fig. 4c-c’) and increased 6-DOT-induced c-Fos in the dHP (Fig. 4d-d’). The same approaches had no effects on dHP activity in naïve (not injected) animals (Supplementary Fig. 8b) and in males (Supplementary Fig. 8c-d’), indicating that VMT-dHP pathway is likely silent in male mice in basal conditions (Supplementary Fig. 8c-c’’, Supplementary Fig. 9b).

We also measured in the *stratum oriens* (*so*) of the dHP the number of total c-Fos^+^ cells and those that colocalized with GAD (Fig. 4e-f), a marker of inhibitory neurons to evaluate any potential change in the inhibitory tone of the dHP in females. The total amount of c-Fos^+^ cells in the *so* was increased, similarly to what we observed in the pyramidal layer (*py*); in parallel, the percentage of colocalizing GAD^+^/c-Fos^+^ cells in *so* decreased after VMT-dHP photoinhibition (Fig. 4f-f’). These data suggested that the inhibition of the VMT-dHP leads to increased hippocampal activity.

VMT-dHP photoinhibition also increased the expression of the eukaryotic translation initiation factor EIF2α, an early regulator of protein synthesis initiation, and decreased the ratio of its phosphorylation (Fig.4g-g’), 1h after the end of the 6-DOT, which can be considered as an index of increased learning-associated protein synthesis^29^. These results indicate that in females, inhibition of VMT-terminals increases glutamatergic transmission in the dHP, possibly by a disinhibitory mechanism, thereby allowing the activation of the LTM molecular processes that males use to allow the expression of spontaneous LTM in conditions of high memory load.

## Discussion

Our study shows that female mice, as well as males^4,8^, express spontaneous memory for up to about 6 different objects at short time delays (1 min). However, keeping this memory load with longer retention intervals (1 to 24 h) unveiled a previously unknown incidental LTM limit in female, but not male mice. Of note, we are not investigating the capacity of LTM, but how the memory load influences memory consolidation during spontaneous encoding. Our spontaneous task reveals sex differences in the engagement dynamics of those circuits, particularly in the LTM manifestation of clear STM traces. Is it that not all the information registered in STM is stabilized into LTM? Studies performed in males may be taken to indicate that high memory load encourages cooperation between STM and LTM mechanisms in the brain^8^; indeed, high memory load favors dHP recruitment in STM tasks. This raises the interesting question of what drives hippocampal recruitment in high memory load tasks? Here, we report a subcortical-to-cortical mechanism that controls the LTM, preventing it in females under high memory load, through VMT hyperactivation, a mechanism which can be exogenously imposed also onto males. Indeed, while males normally recruit the dHP, which favors the expression of LTM traces, females hyperactivate the VMT, which by lowering the dHP activation, limits memory consolidation. The fact that chemogenetic activation of the dHP is sufficient to remove the apparent LTM limit in females strengthens the idea that high memory load somehow promotes cooperation between STM and LTM mechanisms through the engagement of the dHP^8^. In a previous study in males, we reported that increasing the number of objects from 1 to 6 was associated to progressive increase in p-GluR1; however, with 9 objects (an overload condition) p-GluR1 drops to the basal level, despite animals exploring the objects during the study phase, suggesting a lack of recruitment of the dHP. Here we found that the VMT inhibition does not expand memory consolidation beyond its normal limit in males and that the optogenetic inhibition of the VMT-dHP pathway does not change EPSCs in the dHP. Altogether, these findings suggest that females’ memory circuits tend to favor stronger STM, such as retroactive interference resistance, acting through VMT hyperactivation, in contrast to alternative activity that instead favors direct memory consolidation, as in males.

The results from our c-Fos mapping study pointed out the similar recruitment of other cortical regions associated with object processing and memory, such as the ENT and PER and the prefrontal cortices in both sexes, which are not specifically activated for early consolidation of high number of items^30–33^.

Previous evidence in the literature shows that biological relevant stimuli (such as fear), complex tasks and fear memory extinction^34–39^ all lead to VMT activation, which regulates learning and memory consolidation when the interaction between the prefrontal cortex and the dHP is required^40–42^. Therefore, we reasoned that the VMT could represent an important route for transferring information from the STM processing to the HP, which is pivotal for both LTM and high memory load conditions. Our findings add to this body of data, showing that the VMT activation is also determined by information load and that its level of activation is tuned differently in the two sexes. It highlights the importance of memory load as a new, and yet unexplored, factor governing memory consolidation for unique experiences; indeed, we cannot exclude that multiple training session might eliminate or revert the identified sex differences in spontaneous memory.

Our chemogenetic manipulation in the VMT of females and males did not change the total exploration levels in the study phase of the 6-DOT, suggesting that by using this approach we did not affected the general arousal of mice, but we were able to “orchestrate” post-learning VMT activity and consequently dHP engagement and LTM memory capacity. The fact that the degree of VMT activation, which in our study we define as “hyper” vs “hypo” active, is a determinant of the dHP-dependent response is not completely new. Indeed, our evidence is in line with previous data showing that the pattern of stimulation of the VMT is a relevant factor for determining the net effect on dHP-dependent memory also in males: *in vivo* tonic and phasic optogenetic stimulation of the VMT during the acquisition of contextual fear conditioning reduced or augmented fear memory generalization, respectively^43^.

VMT-HP projections are glutamatergic, but by contacting and exciting local GABAergic inhibitory interneurons, VMT projections can indirectly inhibit CA1 pyramidal cell activity. Using extracellular recordings, Dollemen-Van Der Weel et al.^44^ demonstrated that stimulation of the VMT (RE) elicits subthreshold excitatory potentials (no spiking activity) in the dendritic layer of CA1 pyramidal neurons and suprathreshold excitation (spiking activity) in local GABAergic interneurons localized in the *stratum oriens/alveus and radiatum*, which in turn inhibit pyramidal cells. Bertram and Zhang^26^ found a strong excitatory response eliciting population spikes in CA1 when stimulating RE nucleus. As the authors claim this discrepancy is likely due to a stronger stimulation that by recruiting more fibers elicit spiking activity in CA1 pyramidal layer. In a following more detailed study using patch clamp recordings study, Dollemen-Van Der Weel et al. confirmed the strong excitatory connection RE-GABAergic inhibitory neurons^25^. Here we show data from both c-Fos/GAD^+^ neuron mapping and electrophysiological recordings (both *in vivo* and *ex-vivo*) that support the hypothesis that innervation of GABAergic neurons plays a major role in controlling CA1 output (spiking activity). Hence, by silencing VMT inputs, we detected an increased CA1 pyramidal neuron excitation due to disinhibition, caused by a reduced glutamatergic drive onto interneurons, which in turn provide less inhibition to pyramidal cells. This evidence is supported by a recent study^45^, reporting that VMT inputs are exclusively mono-synaptically connected to GABAergic interneurons in the CA1 region, as evidenced with advanced tracing methodology.

From the neurobiological perspective, one possible explanation of the sexually dimorphic action of the VMT on dHP activation, in response to the 6-DOT, is that VMT hyperactivation in females stimulates dHP GABAergic interneurons more than in males. This might be a consequence of the different pattern of VMT stimulation of the dHP, which cannot be distinguished by c-Fos activation alone. Another possibility is that males may have a more resistant dHP activity, as suggested by reports of stronger long-term potentiation and hippocampal theta power activity in males compared to females^46,47^. A higher dHP resistance might in turn limit VMT activation through feedback modulation^27^. These hypotheses are also compatible with the lack of effect of VMT-dHP optogenetic inhibition on EPSCs frequency and amplitude in the dHP of males, as well as on their memory consolidation, in line with previous reports^39,41,42,48,49^.

Although, based on our data, the observed sex differences seem not to be dependent on whether females are in estrous or non-estrous, future studies are needed to evaluate the impact of the level of circulating estrogens on memory performance. Nevertheless, sex-dependent differences in dHP recruitment have already been identified using different behavioral tasks^15^ and a recent paper involving more than 400,000 participants from 38 different countries confirmed this finding^50^, which suggests that these sex differences go beyond the level of circulating estrogens. However, the mechanism underlying the reduced tendency of females to activate the dHP are largely unknown. Most of the evidence indicates a top-down control of memory functions^51,52^. Our data represent a first demonstration of sub-cortical control of dHP recruitment, which might underlie the sex-regulated activation of the dHP, with high potential also for clinical application. Importantly, here we show that the ACC, although activated by exposition to the high memory load request of the 6-DOT, is not functionally involved in early memory consolidation processes that allow the direct transfer of information from the STM to the LTM. This result, still not excluding a role of the ACC in remote high memory load, points back to the importance of the HP in this process and to the brain circuitry that modulates its recruitment. In this the VMT comes to the fore as a key regulator of the cortical mechanisms involved in high load memory consolidation. The negative thalamic regulatory control of dHP activation in high memory load conditions and the consequent limit in the expression of spontaneous memory should not be seen as a structural limit in females, as it may allow them to focus better on ongoing information processing, for instance making them more resistant to retroactive interference. These hypothesis is in line with previous studies showing that thalamic activation sustains attentional control^53^.

Although we do not present direct evidence showing the mechanism governing the sex-different VMT engagement in high memory load, we show that in females there is a spontaneous high load-induced higher VMT recruitment associated to lower hippocampal activation and LTM performance. We cannot exclude the hypothesis that this mechanism was governed by a differential engagement of spatial configurational strategies, in the two sexes, during encoding, which might be activated to lower the information load. However, even if we did not directly explore the latter hypothesis, our experimental design discourages the use of spatially driven object recognition during the test phase, as objects position was changed between the study and the test phase. The fact that post-study phase manipulation of the VMT-dHP pathway removed the LTM limit in female mice, makes highly unlikely the hypothesis that a spatial request in the task has governed the cognitive difference.

The role of thalamic control on cortical activation in memory and cognition is of great current interest. Our data are in line with a recent finding in humans indicating an anti-correlated activation of the mediodorsal thalamus and the hippocampus specifically in females, but not males^54^. We report sex differences in such control mechanisms and demonstrate the existence of these differences in the functional recruitment of the dHP under high memory load, leading in females to limit the expression of spontaneous memory.

The findings that we report suggest a major interaction between subcortical and cortical regions in determining memory processes and are pivotal for elucidating new neurobiological mechanisms of cognitive functions.

## Acknowledgments

Authors would like to thank De Leonibus’ lab members, Dr. Elizabeth Illingworth, Dr. Cornelius Gross, Dr. Vivien Chevaleyre and Prof. Enrico Cherubini for critical revising the manuscript and Dr. Phoebe Ashley-Norman for English editing, and the histology and microscopy facilities of the EMBL (Rome). Authors would also like to thank Miriam Cerullo who helped with VMT-dHP activity correlation and load-dependent activation experiments, Martina Colucci who helped with the immunohistochemistry counting and Caffè Trombetta in via Marsala (Rome) for the logistic support.

## Author contributions

G.T. performed behavioral and immunofluorescence experiments and mice surgeries, conducted all data analysis and statistics, wrote the first draft of the manuscript, reviewed and edited the manuscript, conceptualized optogenetics experiments. V.L. performed behavioral, western blot, immunohistochemistry and immunofluorescence experiments, performed virus localization experiments, conducted data analysis and statistics, produced AAVs for optogenetics, contributed to the review and editing of the manuscript. D.C. and G.S. performed behavioral and immunofluorescence experiments, contributed to data analysis and statistics, contributed to the review and editing of the manuscript. F.E. performed behavioral and immunohistochemistry experiments. A.C. performed and supervised part of the immunofluorescence experiments. M. Gi. contributed to set up optogenetics. M.D.R. performed western blot experiments and produced AAVs for DREADDs, contributed to the review and editing of the manuscript. A.T. contributed to the interpretation of the results and the writing of the manuscript. M.G. conceptualized and performed electrophysiology experiments, contributed to the review and editing of the manuscript. EDL conceived and supervised the study and the experiments, acquired funding, wrote the manuscript. All authors contributed to the discussion and approved the manuscript and supporting information.

## Methods

### Subjects

In this study we used outbred CD1 adult (3 months old) male and female mice (Charles River, Italy) housed in groups of 3-5 subjects.

Housing conditions were maintained at 22 ± 1 °C, with relative humidity of 55 ± 5 %, and with a 12 h light/12 h dark cycle (light: 7:00 am – 7:00 pm). Mice were always tested during the light phase.

All procedures were performed in strict accordance with the European Communities Council directives and Italian laws on animal care (authorization n° 781-2015-PR). Experiments were performed under blind to treatment and sex conditions, unless specified.

### Method for the determination of the estrous cycle phases in female mice

Before behavioral testing, the estrous cycle of females was checked through a visual method^55^, chosen because of its low invasiveness in order to avoid stress to the animals. The validity of the visual inspection method for the determination of the estrous cycle phases was validated before starting the experiments in a subgroup of female mice, as follow: the mouse was gently taken from the resident cage for estrous phase assessment by visual inspection (as described in^55^) and vaginal smears were collected as described in ^56^. Briefly, the tip of an eyedropper filled with distilled water was inserted into the vaginal orifice (less than 1 cm) for flushing, after which one drop of vaginal smear was placed on a slide. Smears were immediately evaluated under a brightfield microscope and representative images were acquired at 20x objective with a Leica Laser Microdissector (Supplementary Fig. 1a). The visual method allows to properly distinguish the estrous from non-estrous (including the metestrus, diestrus and proestrous) with high percentage of concordance, but not each single phase (Supplementary Fig. 1b). Although we found that during estrous female mice also show impaired performance in the 6-DOT at 1 hr delay (Supplementary Fig. 1c), we included in all the other experiments only the non-estrous from the estrous mainly for practical experimental reasons: 1. Estrous is visually easily distinguishable from non-estrous (Supplementary Fig. 1b); 2. Estrous only lasts for 24 hr, which does not fit with experimental designs including 24 h retention interval.

### Habituation procedure

Before each behavioral procedure, mice were first subjected to a habituation procedure. Briefly, the procedure began with manipulation of the mice for 2-5 min. Then, mice were allowed to acclimatize to a personal waiting cage for 10 min. The entire procedure was repeated once a day for at least one week before the beginning of the behavioral procedure.

### Different/Identical Objects Task

The Different/Identical Objects Task (DOT/IOT) was performed as described by Sannino et al^4^. Briefly, animals were isolated for 15 min in a personal waiting cage and then subjected to a familiarization trial (T1; 10 min) in an empty open field (35 × 47 × 60 cm). After 1 min spent in their personal waiting cage, mice were subjected to the study phase (T2; 10 min for the 3, 4 and 6-DOT and 15 min for the 8 and 9-DOT), during which they were allowed to explore the objects until they accumulated 35 sec for the IOT or 35 sec *per* different object for the DOT. Exploration was defined as the time in which the nose of the mouse was in contact with or less than 2 cm from the object ^57^. After a delay of 1 min, for the short-term memory (STM) task, or 1 or 24 h for the long-term memory (LTM) task, animals underwent testing, in which they were exposed to identical copies of the familiar objects and to a new object (T3; 5 min) placed in a randomized position compared to the study phase. Two different types of new objects were alternatively used between animals, and the position of the new object was changed across animals in a random manner. Specifically, we pre-assigned a set of object positions and paired this set between the control animal and the experimental animal, one-by-one (i.e., male vs female or control vs CNO/Light ON), and changed the object set for the successive groups of mice; this allowed us to control for any bias linked to the object used or to its position. Different groups of animals were used for each memory load and delay condition. For c-Fos male/female comparison experiments, total exploration of females was matched to that of males to avoid bias due to different exploration time on the level of c-Fos expression. For this experiment mice were only exposed to the study phase of the 6-DOT.

For DREADDs and optogenetics experiments the same animals were tested two times (15 days apart) and treatments (saline or CNO, light off or light on) were counterbalanced for each test.

The behavior of the animals was recorded by a video-tracking system (Anymaze, Stoelting, USA) and analyzed by a trained observer. We performed off-line behavioral scoring to keep the experimenters blind to the treatments and sex of the animals; when this was not possible, we performed double blind re-scoring performed by another experimenter fully blind on the experiment and obtained comparable scoring results.

Results are expressed as discrimination index (New % Exploration), that is time spent exploring the new object as a percentage of the total exploration time^57^ [(Exploration of New Object/Total object Exploration at T3)*100].

### Retroactive interference procedure

The 6-IOT/DOT was performed as described in the previous paragraph (“Different/Identical Objects Task”). Mice from the interference group were put in the interference cage during the delay between T2 and T3. The interference cage consisted of a standard plastic cage (42.5 × 26.6 × 18.5 cm) filled with small size objects different from those explored in the T2, of which some were embedded with biologically relevant odors (food, sawdust from pups’ cage).

### c-Fos brain mapping by immunohistochemistry

Male and female mice were only exposed to the habituation and the study phase of the 6-DOT (high memory load) and their total object exploration was matched (females to males). Handled naïve sex-matched mice were used as a control group and for normalization of data. All mice were handled as described above.

After 1 h, in order to ensure c-Fos protein expression ^58–60^, mice were deeply anaesthetized and transcardially perfused with 1x PBS followed by 4% paraformaldehyde (PFA; Sigma Aldrich). Brains were collected and post-fixed for 24 h in PFA and passed in a 30% sucrose solution. 30 µm coronal slices were obtained with the use of a vibratome (Leica VT1000 S) and stored in PBS and sodium azide (0.02%) at 4°C until histological processing. 4 to 7 free-floating sections *per* animal were chosen considering the antero-posterior extent for each brain region analysed.

After rinsing in 1x PBS, brain sections were pre-incubated with 0.5 % H_2_O_2_ in 100 % ethanol for 20 minutes to block intrinsic peroxidase activity. Sections were then washed three times with 1x PBS before being pre-incubated for 1 h in a blocking solution made of PBS 0.3% Triton X-100, 1% BSA and 5% normal goat serum (NGS). Blocking solution was then removed and slices were incubated for three days at 4°C with a solution containing 2% NGS, PBS 0.3 % Triton X-100 and 1:400 rabbit anti-c-Fos antibody (sc-52; Santa Cruz Biotechnology). Sections were washed three times in PBS Triton 0.3% (PBS-T) and then incubated for 2 h with anti-rabbit IgG peroxidase-labeled diluted 1:300 in PBS-T containing 1% NGS. Brain sections were exposed to avidin-biotin complex (Avidin/Biotin Blocking Kit; SP-2001, Vector Laboratories, Burlingame, CA). The staining was visualized by color reaction with 3,3′ diaminobenzidinetetrahydrochloride (DAB Peroxidase HRP Substrate Kit, 3,3’-diaminobenzidine; SK-4100; Vector Laboratories, Burlingame, CA) under microscopic control until optimal staining was achieved (approximately 2-5 min). The color reaction was terminated by rinsing sections in Milli-Q water overnight. Finally, sections were mounted on microscope glass slides and coverslipped with Mowiol (4-88; Sigma Aldrich). Control sections, which had not been exposed to primary antibody, were processed in parallel. Images (5x magnification) were taken using the microscope Leica DM6000B with Leica digital camera DFC 480 RGB and Leica application Suite X (LAS X) software.

The number of c-Fos positive cells were manually counted with ImageJ software (National Institute of Health, Bethesda, MD) in all the brain structures for each slice per animal. Number of cells per slice were averaged per group of antero-posterior coordinates and then averaged for each animal. Considering the antero-posterior extent, the resulting values per animal were normalized on naïve average and expressed as percentage of naïve.

### Preparation of AAVs

Plasmids for DREADDs and optogenetic experiments were purchased from Addgene.

For DREADDs experiments we used pAAV-CaMKIIa-HA-rM3D(Gs)-IRES-mCitrine (#50468) and pAAV-CaMKIIa-HA-hM4D(Gi)-IRES-mCitrine (#50467).

For optogenetic experiments we used pAAV-hSyn-eNpHR 3.0-EYFP (#26972).

Plasmids were amplified and purified using EndoFree Plasmid Mega Kit (Qiagen), following manufacturer’s instruction and AAVs were assembled by the TIGEM AAV Vector Core.

AAV serotypes produced resulted in the following titer (GC/ml): 1.4 × 10^12^ for AAV 2/5-CaMKIIa-HA-rM3D(Gs)-IRES-mCitrine, 2.1 × 10^12^ for AAV 2/5-CaMKIIa-HA-hM4D(Gi)-IRES-mCitrine and 8.4 × 10^12^ for AAV 2/2-h-SYN-eNpHR3.0-EYFP.

### Stereotaxic surgery

For virus injection, mice were anaesthetized with a mixture of tiletamine/zolazepam (Zoletil; 25 mg/Kg) and xylazine (5 mg/Kg) and placed on a stereotaxic apparatus (Stoelting, USA). The skull was exposed and small craniotomies were made at the following coordinates: antero-posterior (AP) −0.82 mm and mediolateral (ML) ± 0.2 mm from bregma, and dorso-ventral (DV) −4.3 mm from the dura for the ventral midline thalamus (VMT); AP – 1.80 mm, ML ± 1.5 mm from bregma and DV −1.6 mm from the dura for the dorsal hippocampus (dHP); AP + 0.86 mm, ML ± 0 mm from bregma and DV −1.75 mm from the dura for the anterior cingulate cortex (ACC). Injection volume was 0.4 μL for the VMT and ACC and 0.5 μL per side for the HP.

For fiber optic implantation, during the same surgery one or two 200 μm core (0.39 NA) optic fibers were implanted at the coordinates AP – 0.82 mm, ML ± 0.2 mm from bregma and DV −4.1 mm from the dura for the VMT, and AP – 1.80 mm, ML ± 1.6 mm from bregma and DV −1.5 mm from the dura for the HP. Two screws were fixed on the skull to ensure implant stability. Optic fibers were fixed in place with dental cement (Meliodent, Heraeus).

For FluoroGold (FG) injection female mice were anaesthetized as before and 0.5 μL of a 4% FG solution dissolved in saline (NaCl 0.9%) was unilaterally injected in the HP at the same coordinates used for the viral injection.

At the end of the surgical procedure mice were intraperitoneally (ip) injected with 1 ml saline solution (NaCl 0.9%) and left to recover for 10-15 days.

### *In vivo* drug injection for DREADDs experiments

Two weeks after surgery mice underwent behavioral testing. 30 min before the beginning of the familiarization trial (T1) or immediately after the end of the training phase (T2), mice were injected with vehicle (Veh; NaCl 0,9%) or CNO (1 mg/kg; Hello Bio). At the end of study phase, mice were left to rest in their waiting cages for 1 h, after which they were put back in the resident cage. 24 h later mice underwent to the test phase.

### Light stimulation protocol for optogenetics experiments

Four to five weeks post-AAV injection, mice underwent behavioral testing and light stimulation protocol. Immediately after the end of the study phase the implantable fibers were coupled to a 120-mW, 532-nm diode-pumped solid-state laser (Laserlands, China) using a one-input one-output rotary joint (Thorlabs, USA) connected to a bifurcated fiber bundle (Thorlabs, USA). Power output from the tip of the bifurcated fiber bundle was about 11-12 mW. Immediately, after the end of the study phase 30 min of continuous light stimulation (light-on) or no light (light off) was delivered. After photostimulation mice were left to rest for an additional 30 min in their waiting cages after which they were put back in the resident cage for the night. 24 h later mice underwent 5 min test phase.

### AAVs and FluoroGold expression verification

At the end of the behavioral procedure for DREADDs experiment, mice were deeply anaesthetized and transcardially perfused with PBS followed by 4% paraformaldehyde (PFA; Sigma Aldrich). Brains were dissected and post-fixed for 24 h in PFA 4%, then they were washed in PBS and put in 30% sucrose solution. 30 µm coronal slices were obtained with the use of a cryostat (Leica CM3050 S) and stored in PBS and sodium azide 0.02% at 4° C until the beginning of the procedure. Several free-floating sections *per* mouse were chosen for the HP and the VMT to evaluate the placement and extension of AAV injection.

For DREADDs experiments, after rinsing in 1x PBS, brain sections were incubated in blocking solution made of NGS 5% and 0.3% Triton X-100 in PBS for 1 h. Blocking solution was then removed and replaced with primary antibody solution made of NGS 1%, Triton X-100 0.3% and rabbit anti-HA tag (#3724; Cell Signaling) diluted 1:500 overnight at 4°C. Sections were washed three times in 1x PBS and then incubated for 2 h at room temperature with secondary antibody solution made with NGS 1%, Triton X-100 0.3% and goat-anti-rabbit Alexa Fluor 647 (Merck Millipore) diluted 1:300 in 1x PBS. After three washes with 1x PBS, sections were incubated for 10 mins with 4′,6-diamidino-2-phenylindole (DAPI), for nucleic acid staining, at room temperature and then washed three times with 1x PBS. Finally, sections were mounted on microscope glass slides and coverslipped with Mowiol (4-88; Sigma Aldrich). Control sections, which had not been exposed to primary antibody, were processed in parallel.

Slices were manually analyzed under a Nikon Ni-E fluorescence microscope equipped with Nikon DS-Ri2 camera and NIS-Elements C 4.20 (Nikon, Italy). Only mice expressing the reporter HA in the HP or in the VMT were included in the statistical analysis.

For the optogenetic experiment, the brain sampling protocol was the same as that described previously for DREADDs, but in this case, HP and VMT sections were only incubated for 10 mins with DAPI and subsequently mounted on microscope glass slides and coverslipped with Mowiol.

Slices were manually analyzed and acquired under a Nikon Ni-E fluorescence microscope equipped with Nikon DS-Ri2 camera and NIS-Elements C 4.20 (Nikon, Italy). Only mice expressing the reporter EYFP in the VMT and the HP were included in the statistical analysis.

FluoroGold (FG) injected mice were deeply anaesthetized, transcardially perfused and their brains were treated as described above. FG labelled cells were visualized on 40 µm coronal slices counterstained with a 5 μg/ml propidium iodide (Santa Cruz Biotechnology) staining solution in PBS 1x for 10 min. Mosaic reconstruction of dHP and VMT slices were acquired with a 6x objective, while VMT magnification - showing the nucleus reuniens - was acquired with a 10x NA 0.2 objective with a Leica Laser Microdissector.

### *Ex vivo* electrophysiological recordings in slices

Transverse hippocampal slices (320 μm tick) were obtained using a standard protocol^61^ from 3 months old mice injected 4-5 weeks before with AAV-hSyn-eNpHR 3.0-EYFP in the VMT. Briefly, after being anesthetized with an intraperitoneal injection of a mixture of tiletamine/zolazepam (80mg/Kg) and xylazine (10 mg/Kg), mice were decapitated. The brain was quickly removed from the skull, placed in artificial cerebrospinal fluid (ACSF) containing (in mM): sucrose 75, NaCl 87, KCl 2.5, NaH_2_PO_4_ 1.25, MgCl_2_ 7, CaCl_2_ 0.5, NaHCO_3_ 25, glucose 25. After recovery, an individual slice was transferred to a submerged recording chamber and continuously perfused at room temperature with oxygenated ACSF at a rate of 3 ml/min. ASCF saturated with 95% O_2_ and 5% CO_2_ and contained in mM: NaCl 125, KCl 2.5, NaH_2_PO_4_ 1.25, MgCl_2_ 1, CaCl_2_ 2, NaHCO_3_ 25, glucose 10.

Cells were visualized with a 60X water immersed objective mounted on an upright microscope (Nikon, eclipse FN1) equipped with a CCD camera (Scientifica, UK). eNHpR+ fibers were photoinhibited with continuous (5-10 minutes) green light delivered through the objective and generated by a 535 nm LED (pE2, CoolLED, UK) under the control of the digital output of the amplifier. Whole-cell patch clamp recordings, in voltage and current clamp modes, were performed with a MultiClamp 700B amplifier (Axon Instruments, Sunnyvale, CA, USA). Patch electrodes were pulled from borosilicate glass capillaries (WPI, Florida, US); they had a resistance of 3-4 MΩ when filled with an intracellular: K gluconate 127, KCl 6, HEPES 10, EGTA 1, MgCl_2_ 2, MgATP 4, MgGTP 0.3; the pH was adjusted to 7.2 with KOH; the osmolarity was 290–300 mOsm. Spontaneous postsynaptic excitatory currents (EPSCs) were recorded at the equilibrium potential for chloride (E_Cl-_) that was approximately −65 mV based on the Nernst equation. Membrane potential values were not corrected for liquid junction potentials. The stability of the patch was checked by repetitively monitoring the input and series resistance during the experiments. Series resistance (10-20 MΩ) was not compensated. Cells exhibiting 15% changes were excluded from the analysis.

Data were transferred to a computer hard disk after digitization with an A/D converter (Digidata 1550, Molecular Devices, Sunnyvale, CA, USA). Data acquisition (digitized at 10 kHz and filtered at 3 kHz) was performed with pClamp 10.4 software (Molecular Devices, Sunnyvale, CA, USA). Input resistance and cells capacitance were measured online with the membrane test feature of the pClamp software. Spontaneous EPSCs were analyzed with pClamp 10.4 (Molecular Devices, Sunnyvale, CA, USA). This program uses a detection algorithm based on a sliding template. The template did not induce any bias in the sampling of events because it was moved along the data trace one point at a time and was optimally scaled to fit the data at each position.

### *In vivo* electrophysiological recordings from anesthetized animals

Mice injected 4-5 weeks before with AAV-hSyn-eNpHR 3.0-EYFP in the VMT were anesthetized with i.p. injection of a mixture of tiletamine/zolazepam (Zoletil; 80mg/Kg) and xylazine (10 mg/Kg) to induce anesthesia before surgery and during recordings. Temperature was maintained between 36–37°C using a feedback-controlled heating pad (FHC). A craniotomy for stimulation and recording sites was drilled between 1.6 and 2.0 mm posterior from bregma, and lateral coordinates were adjusted after extracellular mapping to locate the dorsal CA1 pyramidal cell layer. Light activation of eNHpR+ fibers was achieved with 50 mW 532 nm laser (SLOC Lasers, Shangai) delivered through an optical fiber (200 µm diameter, 0.22 numerical aperture NA) lowered with 10° angle in the *stratum radiatum*. Power output at the tip of the fiber was 3-4 mW. Continuous light was triggered for 5-10 minutes.

Extracellular recordings of local field potentials from CA1 were obtained with glass electrodes (Hingelberg, Malsfeld, Germany) prepared with a vertical puller PP-830 (Narishige), and the tip was broken to obtain a resistance of 1-2 MΩ. The distance between the optical fiber and the recording electrode was 150-200 µm. Electrodes were filled with a standard Ringer’s solution containing the following (in mM): NaCl 135, KCl 5.4, HEPES 5, CaCl_2_ 1.8, and MgCl_2_ 1.

Recordings were obtained with a Multiclamp 700B amplifier connected to the Digidata 1550 system. Data were acquired (digitized at 10 kHz and filtered at 3 kHz) with pClamp 10.4 software and analyzed off-line with Clampfit 10.4 (Molecular Devices).

Traces were high pass filtered (300 Hz) and a clampfit algorithm was used to detect extracellular spikes (MUA) setting a threshold of three standard deviations from the noise. The spontaneous MUA was estimated during 5-10 min before, during and after light stimulation.

### Western blot

At the end of the DREADDs behavioral procedure, after one week, mice were injected again with CNO or Veh and, after 65 min, they were sacrificed through cervical dislocation and their brains were collected. The dorsal hippocampi (HP) were dissected using a brain matrix (Kopf Instruments). GluA1 post-translational mechanisms in the hippocampus are not constitutively activated or inhibited, rather they are induced by exposure to the 6-DOT^8^. Consequently, in the experiment reported in Fig. 3a, we expected that chemogenetic activation of the dHP in females would be sufficient to induce an increase in S845 phosphorylation in females, as compared to basal conditions (saline injected). In contrast, in Fig. 3d, chemogenetic activation of VMT in males was expected to cause a decrease of S845 activation in the dHP, which could have been difficult to detect in basal conditions (as also evidenced by the lack of effects of optogenetic inhibition of the VMT-dHP pathway in Supplementary Fig. 8c-c’), without any stimulus; therefore, we challenged male mice with the 6-DOT, to promote S845 phosphorylation, while preventing it in the CNO group, by hyperactivating the VMT; this allowed us to detect a decrease in S845 in male mice.

HP sections were homogenized with homogenizing buffer composed of 0.32 M sucrose, 1 mM EDTA, 1 mg/mL BSA, 5 mM HEPES pH 7.4, 1 mM PMSF, 2 mM sodium orthovanadate, 10 mM sodium fluoride, 1x Sigma protease inhibitor cocktail in distilled H_2_O. Protein concentration was determined by Bradford assay (Bio-Rad). Dorsal HP homogenates aliquots (20 µg/lane) from CNO and saline treated mice were first boiled at 95°C for 5 min and then separated on 10% polyacrylamide gel and transferred on PVDF membranes (Immobilon-P transfer membrane; Merck Millipore). Membranes were then blocked with 5% non-fat powdered milk in TBS + 0.01% Tween-20 (TBS-T) for 1 h at room temperature followed by overnight incubation at 4°C with rabbit anti-glutamate receptor 1 (AMPA subtype) phosphoSer 845 antibody (1:500; ab76321, Abcam), rabbit anti-glutamate receptor 1 (AMPA subtype) antibody (1:500; ab31232, Abcam) and anti-β-actin (1:5000, MAB1501, Millipore).

Subsequently, blots were washed with TBS-T and secondary antibodies (1:5000; Bio-Rad) were applied for 1 h at room temperature. Bands were detected by chemiluminescence and quantified using ImageJ software (NIH, Bethsda, MD). Data are expressed as % from Veh treated samples.

We reported normalized data to the Vehicle group because we calculated the percentage relative to the control for each blot, to correct for any type of difference relative to different gels (for example recycling of primary antibody or time to ECL exposure). Uncropped and unprocessed scans of the blots are provided in the Source Data file.

### c-Fos counting and GAD67 co-localization by immunofluorescence

After one week from the behavioral evaluation, mice from the VMT-HP optogenetic inhibition experiment were exposed to the habituation and the study phase of the 6-DOT (high memory load) followed by 30/30 min of light on/off stimulation or 60 min light off as previously done. Mice were then deeply anaesthetized and transcardially perfused with 1x PBS followed by 4% paraformaldehyde (PFA; Sigma Aldrich). Brains were collected and post-fixed for 24 h in PFA and passed in a 30% sucrose solution. 30 µm coronal slices were obtained with the use of a vibratome (Leica VT1000 S) and stored in PBS and sodium azide (0.02%) at 4°C until histological processing. Slices were incubated with primary antibodies against: c-Fos (1:1000, 226 017, Synaptic System) and GAD67 (1:700, MAB5406, Merck Millipore) after a blocking step with PBS-Triton X-100 0.3% and 5% normal goat serum (NGS). Slices were subsequently washed with PBS and incubated for 1 h with the proper secondary antibodies (1:400, goat anti-rat Alexa-Fluor® 568, ab175476, abcam; 1:400, goat anti-mouse Alexa-Fluor® 488, AP124JA4, Merck Millipore). For nuclear counterstaining slices were incubated with DAPI (1:1000, 10 min, D1306, Invitrogen) before mounting and inclusion with a Mowiol 4-88 (Sigma Aldrich) solution.

Images of c-Fos and GAD67 co-localization were acquired as z-stacks (9 sections every 2 μm) using a spinning disk confocal microscope (X-light V3 Crest Optics) using a 25x NA 1.05 silicon oil immersion objective. All images were acquired using a Prime BSI with 2048 × 2048 pixels. For all the experiments 3-5 slice *per* mouse and 3-4 mice *per* group were acquired and analyzed using QuPath 0.2.3^62^. For each slice three alternated z-stack were sampled with a region of interest (ROI) in the pyramidal layer (*py*) and in the stratum oriens (*so*) of the CA1. c-Fos^+^ cells were first counted in the *py* and the *so* independently with the cell detection tool by QuPath. Subsequently, the c-Fos^+^ cells of the *so* were classified as GAD^+^ or GAD^-^ using the single measurement classifier tool by QuPath. c-Fos^+^ and c-Fos^+^/GAD^+^ cells were then summed per area per ROI for each HP side and averaged per slice. The percentage of c-Fos^+^/GAD^+^ cells was calculated for the total number of c-Fos^+^ cells. Slices were averaged *per* groups (light off *vs* light on) for female and male mice.

### p-EIF2α/EIF2α quantification in excitatory neurons

One week following the behavioral evaluation, mice from the VMT-HP optogenetic inhibition experiment were exposed to the habituation and study phase of the 6-DOT (high memory load) followed by 30/30 min of light on/off stimulation or 60 min light off as previously done. Mice were then deeply anaesthetized and transcardially perfused with 1x PBS followed by 4% paraformaldehyde (PFA; Sigma Aldrich). Brains were collected and post-fixed for 24 h in PFA and passed in a 30% sucrose solution. 30 µm coronal slices were obtained with the use of a vibratome (Leica VT1000 S) and stored in PBS and sodium azide (0.02%) at 4°C until histological processing. Slices were incubated with primary antibodies against: p-EIF2α (Ser51) (1:200, 3597, Cell Signaling) or EIF2α (1:800, 5324, Cell Signaling), and GAD67 (1:700, MAB5406, Merck Millipore) and NeuN (1:500, ABN90, Merck Millipore) after a blocking step with PBS-Triton X-100 0.3% and 5% normal goat serum (NGS). Slices were subsequently washed with PBS and incubated for 2 h with the proper secondary antibodies (1:300, goat anti-rabbit Alexa-Fluor® 647, AP187SA6, Merck Millipore; 1:300, goat anti-guinea pig Alexa-Fluor® 568, ab175714, Abcam; 1:400, goat anti-mouse Alexa-Fluor® 488, AP124JA4, Merck Millipore). For nuclear counterstaining slices were incubated with DAPI (1:1000, 10 min, D1306, Invitrogen) before mounting and inclusion with Mowiol 4-88 (Sigma Aldrich) solution.

Images of p-EIF2α/GAD67/NeuN and EIF2α/GAD67/NeuN were acquired using a confocal microscope (Leica SP5) with a 40x objective and a 1.5x digital zoom. All images were acquired with a 1024×1024 pixel resolution. For all the experiments 3 slice *per* mouse and 3 mice *per* group were acquired and analyzed using ImageJ (NIH). Each image was sampled in the GAD67^+^ channel with 4 regions of interest (ROIs) positioned on the excitatory neurons, using the pattern of GAD67 cytoplasmatic expression as a discriminatory marker between excitatory neurons (GAD67^-^). NeuN was used as a counterstaining to recognize neurons. The fluorescence intensity (AU) of p-EIF2α or EIF2α was measured in each ROI and averaged for category of excitatory neurons (GAD67^-^) *per* groups (light off *vs* light on) for female and male mice. The number of p-EIF2α^+^ spots was correlated to the fluorescence intensity values per ROI for a subgroup of slice from males and females to prove the reliability of the fluorescence intensity signal as an index of the amount of p-EIF2α levels (Supplementary Fig. 9c). Fluorescence intensity was averaged per slices of each experimental condition.

### Statistics

Data were analyzed with the software: Statview 5.0, Statistica 7, GraphPad Prism 8 and G*power 3.1. The significance level was set at p < 0.05 for all the experiments; data are expressed as mean ± standard errors (SEM). The number of mice per group was calculated *a priori* with power analysis using G^*^Power 3.1 software^63–65^ with *α* = 0.05 and power (1 − *β*) = 0.80, and data were inspected for normal distribution through a Shapiro-Wilk test. Significant outliers were calculated at the alpha level 0.05, through the online free software Outlier Calculator by GraphPad (available at https://www.graphpad.com/quickcalcs/Grubbs1.cfm), which employs Grubbs’ test to define a significant outlier.

Discrimination index at T3 (test phase) and other two-levels variables were analyzed with a one-way or a three-way ANOVA or a Mann-Whitney test in case normal distribution was not respected. Depending on the experiment, sex (two levels: male, female), load (two levels:6-IOT, 6-DOT), delay (two levels: 1 min, 24 h) and treatment (two levels: Veh, CNO for DREADDs experiments, or light on, light off for optogenetics experiments) were used as between factors.

For single object comparisons, we used a one-way ANOVA for repeated measures with objects (six levels: New, F1-F5) as repeated measures, followed by Dunnett post-hoc analysis only when the ANOVA was significant. Counting of c-Fos^+^ cells per slice was averaged per animal and, for each brain region, data were normalized on naïve average considering the antero-posterior extent, thus values were expressed as % on naïve expression. A two-way ANOVA for sex (two levels: male, female) and test (two levels: naïve, test) was used. Bonferroni post-hoc analysis was carried out when the p-value for interaction of sex x test was significant; if this was not the case a one-way ANOVA for sex or treatment separately was used.

For Western Blot analysis, a one-way ANOVA for treatment (two levels: saline, CNO) was used to analyze data for GluR1 p-S845 and data, normalized on GluR1 levels, are expressed as % vs 3 months treated samples.

For MUA analysis Wilcoxon matched-pairs signed rank test was used to assess CNO effect on firing rate.

## Data availability statement

All data is available upon reasonable request to the corresponding author. Source data underlying the main and supplementary figures are provided as a Source Data file.

## References

1. Frankland, P. W. & Bontempi, B. The organization of recent and remote memories. Nat. Rev. Neurosci. 6, 119–30 (2005).

2. Lee, I., Hunsaker, M. R. & Kesner, R. P. The role of hippocampal subregions in detecting spatial novelty. Behav. Neurosci. 119, 145–153 (2005).

3. Save, E., Poucet, B., Foreman, N. & Buhot, M. C. Object Exploration and Reactions to Spatial and Nonspatial Changes in Hooded Rats Following Damage to Parietal Cortex or Hippocampal Formation. Behav. Neurosci. 106, 447–456 (1992).

4. Sannino, S. et al. Role of the dorsal hippocampus in object memory load. Learn. Mem. 19, 211–218 (2012).

5. Sugita, M., Yamada, K., Iguchi, N. & Ichitani, Y. Hippocampal NMDA receptors are involved in rats’ spontaneous object recognition only under high memory load condition. Brain Res. 1624, 370–379 (2015).

6. Rodo, C., Sargolini, F. & Save, E. Processing of spatial and non-spatial information in rats with lesions of the medial and lateral entorhinal cortex: Environmental complexity matters. Behav. Brain Res. 320, 200–209 (2017).

7. Iemolo, A. et al. Reelin deficiency contributes to long-term behavioral abnormalities induced by chronic adolescent exposure to Δ9-tetrahydrocannabinol in mice. Neuropharmacology 187, (2021).

8. Olivito, L. et al. Phosphorylation of the AMPA receptor GluA1 subunit regulates memory load capacity. Brain Struct. Funct. 221, 591–603 (2016).

9. Beason-Held, L. L., Rosene, D. L., Killiany, R. J. & Moss, M. B. Hippocampal formation lesions produce memory impairment in the rhesus monkey. Hippocampus 9, 562–74 (1999).

10. Levy, D. A., Hopkins, R. O. & Squire, L. R. Impaired visual and odor recognition memory in patients with hippocampal lesions. Learn. Mem. 11, 794–796 (2003).

11. Levy, D. A., Hopkins, R. O. & Squire, L. R. Impaired odor recognition memory in patients with hippocampal lesions. Learn. Mem. 11, 794–6 (2004).

12. Spets, D. S., Jeye, B. M. & Slotnick, S. D. Different patterns of cortical activity in females and males during spatial long-term memory. Neuroimage 199, 626–634 (2019).

13. Spets, D. S. & Slotnick, S. D. Are there sex differences in brain activity during long-term memory? A systematic review and fMRI activation likelihood estimation meta-analysis. Cogn. Neurosci. 00, 1–11 (2020).

14. Cimadevilla, J. M., Conejo, N. M., Miranda, R. & Arias, J. L. Sex differences in the Morris water maze in young rats: Temporal dimensions. Psicothema 16, 611–614 (2004).

15. Torromino, G., Maggi, A. & Leonibus, E. De. Estrogen-dependent hippocampal wiring as a risk factor for age-related dementia in women. Prog. Neurobiol. (2020). doi:10.1016/j.pneurobio.2020.101895

16. Spiers, H. J., Coutrot, A. & Hornberger, M. Explaining World-Wide Variation in Navigation Abilityfrom Millions of People: Citizen Science ProjectSea Hero Quest. Top. Cogn. Sci. 0, 1–19 (2021).

17. Coutrot, A. et al. Virtual navigation tested on a mobile app is predictive of real-world wayfinding navigation performance. PLoS One 14, 1–15 (2019).

18. Juhas, I., Baĉanac, L. & Kozoderović, J. The Most Common Errors in Orienteering and Their Relation To Gender, Age and Competition Experience. Facta Univ. Ser. Phys. Educ. Sport 14, 211–226 (2016).

19. Olivito, L., De Risi, M., Russo, F. & De Leonibus, E. Effects of pharmacological inhibition of dopamine receptors on memory load capacity. Behav. Brain Res. 359, 197–205 (2019).

20. Shrager, Y., Levy, D. A., Hopkins, R. O. & Squire, L. R. Working Memory and the Organization of Brain Systems. 28, 4818–4822 (2008).

21. Aggleton, J. P. & Brown, M. W. Episodic memory, amnesia, and the hippocampal-anterior thalamic axis. Behav. Brain Sci. 22, 425–444; discussion 444-489 (1999).

22. Kesner, R. P., Ravindranathan, A., Jackson, P., Giles, R. & Chiba, A. A. A neural circuit analysis of visual recognition memory: Role of perirhinal, medial, and lateral entorhinal cortex. Learn. Mem. 8, 87–95 (2001).

23. Wilson, D. I. G. et al. Lateral entorhinal cortex is critical for novel object-context recognition. Hippocampus 23, 352–366 (2013).

24. Dolleman-Van Der Weel, M. J. & Witter, M. P. Projections From the Nucleus Reuniens Thalami to the Entorhinal Cortex, Hippocampal Field CA1, and the Subiculum in the Rat Arise From Different Populations of Neurons. J. Comp. Neurol. 364537450, (1996).

25. Dolleman-van DerWeel, M. J. & Witter, M. P. Nucleus reuniens thalami innervates g aminobutyric acid positive cells in hippocampal field CA1 of the rat. Neurosci. Lett. 278, 145–148 (2000).

26. Bertram, E. H. & Zhang, D. X. Thalamic excitation of hippocampal CA1 neurons: A comparison with the effects of CA3 stimulation. Neuroscience 92, 15–26 (1999).

27. Dolleman-van DerWeel, M. J. et al. The nucleus reuniens of the thalamus sits at the nexus of a hippocampus and medial prefrontal cortex circuit enabling memory and behavior. Learn. Mem. 26, 191–205 (2019).

28. Roth, B. L. DREADDs for Neuroscientists. Neuron 89, 683–694 (2016).

29. Sharma, V. et al. eIF2α controls memory consolidation via excitatory and somatostatin neurons. Nature (2020). doi:10.1038/s41586-020-2805-8

30. Ennaceur, A., Neave, N. & Aggleton, J. P. Spontaneous object recognition and object location memory in rats: The effects of lesions in the cingulate cortices, the medial prefrontal cortex, the cingulum bundle and the fornix. Exp. Brain Res. 113, 509–519 (1997).

31. Reid, J. M., Jacklin, D. L. & Winters, B. D. Delineating prefrontal cortex region contributions to crossmodal object recognition in rats. Cereb. Cortex 24, 2108–2119 (2014).

32. Pezze, M. A., Marshall, H. J., Fone, K. C. F. & Cassaday, H. J. Dopamine D1 receptor stimulation modulates the formation and retrieval of novel object recognition memory: Role of the prelimbic cortex. Eur. Neuropsychopharmacol. 25, 2145–2156 (2015).

33. Pezze, M. A., Marshall, H. J., Fone, K. C. F. & Cassaday, H. J. Role of the anterior cingulate cortex in the retrieval of novel object recognition memory after a long delay. 310–317 (2017).

34. Ramanathan, K. R., Ressler, R. L., Jin, J. & Maren, S. Nucleus reuniens is required for encoding and retrieving precise, hippocampal-dependent contextual fear memories in rats. J. Neurosci. 38, 9925–9933 (2018).

35. Silva, B. A., Burns, A. M. & Gräff, J. A cFos activation map of remote fear memory attenuation. Psychopharmacology (Berl). 236, 369–381 (2019).

36. Silva, B. A. et al. A thalamo-amygdalar circuit underlying the extinction of remote fear memories. Nat. Neurosci. (2021). doi:10.1038/s41593-021-00856-y

37. Lin, Y. J., Chiou, R. J. & Chang, C. H. The reuniens and rhomboid nuclei are required for acquisition of pavlovian trace fear conditioning in rats. eNeuro 7, 1–15 (2020).

38. Loureiro, M. et al. The ventral midline thalamus (Reuniens and rhomboid nuclei) contributes to the persistence of spatial memory in rats. J. Neurosci. 32, 9947–9959 (2012).

39. Barker, G. R. I. & Warburton, E. C. A critical role for the nucleus reuniens in long-term, but not short-term associative recognition memory formation. J. Neurosci. 38, 3208–3217 (2018).

40. Ito, H. T., Zhang, S., Witter, M. P., Moser, E. I. & Moser, M. A prefrontal–thalamo–hippocampal circuit for goal-directed spatial navigation. Nature 522, (2015).

41. Cholvin, T., Hok, X. V., Giorgi, L., Chaillan, F. A. & Poucet, X. B. Ventral Midline Thalamus Is Necessary for Hippocampal Place Field Stability and Cell Firing Modulation. J. Neurosci. 38, 158–172 (2018).

42. Hauer, B. E., Pagliardini, S. & Dickson, C. T. The reuniens nucleus of the thalamus has an essential role in coordinating slow-wave activity between neocortex and hippocampus. eNeuro 6, (2019).

43. Xu, W. & Südhof, T. C. A Neural Circuit for Memory Specificity and Generalization. Science (80-.). 339, 1290– 1296 (2013).

44. Weel, M. J. D. Der, Silva, F. H. L. & Witter, M. P. Nucleus Reuniens Thalami Modulates Activity in Hippocampal Field CA1 through Excitatory and Inhibitory Mechanisms. 17, 5640–5650 (1997).

45. Andrianova, L. et al. Hippocampal CA1 pyramidal cells do not receive monosynaptic input from thalamic nucleus reuniens. bioRxiv 2021.09.30.462517 (2021).

46. Maren, S., De Oca, B. & Fanselow, M. S. Sex differences in hippocampal long-term potentiation (LTP) and Pavlovian fear conditioning in rats: positive correlation between LTP and contextual learning. Brain Res. 661, 25– 34 (1994).

47. Juárez, J. & Corsi-Cabrera, M. Sex differences in interhemispheric correlation and spectral power of EEG activity. Brain Res. Bull. 38, 149–151 (1995).

48. Ito, H. T., Moser, E. I. & Moser, M. Supramammillary Nucleus Modulates Spike-Time Coordination in the Prefrontal-Thalamo-Hippocampal Circuit during Navigation Article Supramammillary Nucleus Modulates Spike-Time Coordination in the Prefrontal-Thalamo-Hippocampal Circuit during Navigation. Neuron 99, 576-587.e5 (2018).

49. Dolleman-Van Der Weel, M. J., Morris, R. G. M. & Witter, M. P. Neurotoxic lesions of the thalamic reuniens or mediodorsal nucleus in rats affect non-mnemonic aspects of watermaze learning. Brain Struct. Funct. 213, 329–342 (2009).

50. Coutrot, A. et al. Entropy of city street networks linked to future spatial navigation ability. Nature 604, 104–110 (2022).

51. Torromino, G. et al. Offline ventral subiculum-ventral striatum serial communication is required for spatial memory consolidation. Nat. Commun. 10, 1–9 (2019).

52. Vetere, G. et al. An inhibitory hippocampal–thalamic pathway modulates remote memory retrieval. Nat. Neurosci. 24, 685–693 (2021).

53. Schmitt, L. I. et al. Thalamic amplification of cortical connectivity sustains attentional control. Nat. Publ. Gr. 545, (2017).

54. Spets, D. S. & Slotnick, S. D. Thalamic Functional Connectivity during Spatial Long-Term Memory and the Role of Sex. Brain Sci. 10, 1–15 (2020).

55. Byers, S. L., Wiles, M. V, Dunn, S. L. & Taft, R. A. Mouse Estrous Cycle Identification Tool and Images. 7, 2–6 (2012).

56. Goldman, J. M., Murr, A. S. & Cooper, R. L. The rodent estrous cycle: Characterization of vaginal cytology and its utility in toxicological studies. Birth Defects Res. Part B - Dev. Reprod. Toxicol. 80, 84–97 (2007).

57. Ainge, J. A. et al. The role of the hippocampus in object recognition in rats : Examination of the influence of task parameters and lesion size. Behav. Brain Res. 167, 183–195 (2006).

58. Guzowski, J. F., Setlow, B., Wagner, E. K. & McGaugh, J. L. Experience-dependent gene expression in the rat hippocampus after spatial learning: a comparison of the immediate-early genes Arc, c-fos, and zif268. J. Neurosci. 21, 5089–5098 (2001).

59. Maviel, T., Durkin, T. P., Menzaghi, F. & Bontempi, B. Sites of Neocortical Reorganization Critical for Remote Spatial Memory. Science (80-.). 305, 96–99 (2004).

60. De Leonibus, E., Mele, A., Oliverio, A. & Pert, A. Distinct pattern of c-fos mRNA expression after systemic and intra-accumbens amphetamine and MK-801. Neuroscience 115, 67–78 (2002).

61. Bischofberger, J., Engel, D., Li, L. & Al., E. Patch-clamp recording from mossy fiber terminals in hippocampal slices. Nat Protoc 1, 2075–2081 (2006).

62. Bankhead, P. et al. QuPath: Open source software for digital pathology image analysis. Sci. Rep. 7, 1–7 (2017).

63. Faul, F., Erdfelder, E., Buchner, A. & Lang, A. Statistical power analyses using G*Power 3.1: tests for correlation and regression analyses. Behav Res Methods 41, 1149–60 (2009).

64. Capuozzo, A. et al. Fluoxetine ameliorates mucopolysaccharidosis type IIIA. Mol Ther. S1525-0016, (2022).

65. Sonzogni, M. et al. A behavioral test battery for mouse models of Angelman syndrome: A powerful tool for testing drugs and novel Ube3a mutants. Mol. Autism 9, 1–19 (2018).

