## Supplementary Information for "Thalamo-hippocampal pathway regulates incidental memory load in mice"

**This PDF file includes:**

Supplementary Figures 1-9
Supplementary Tables 1-5

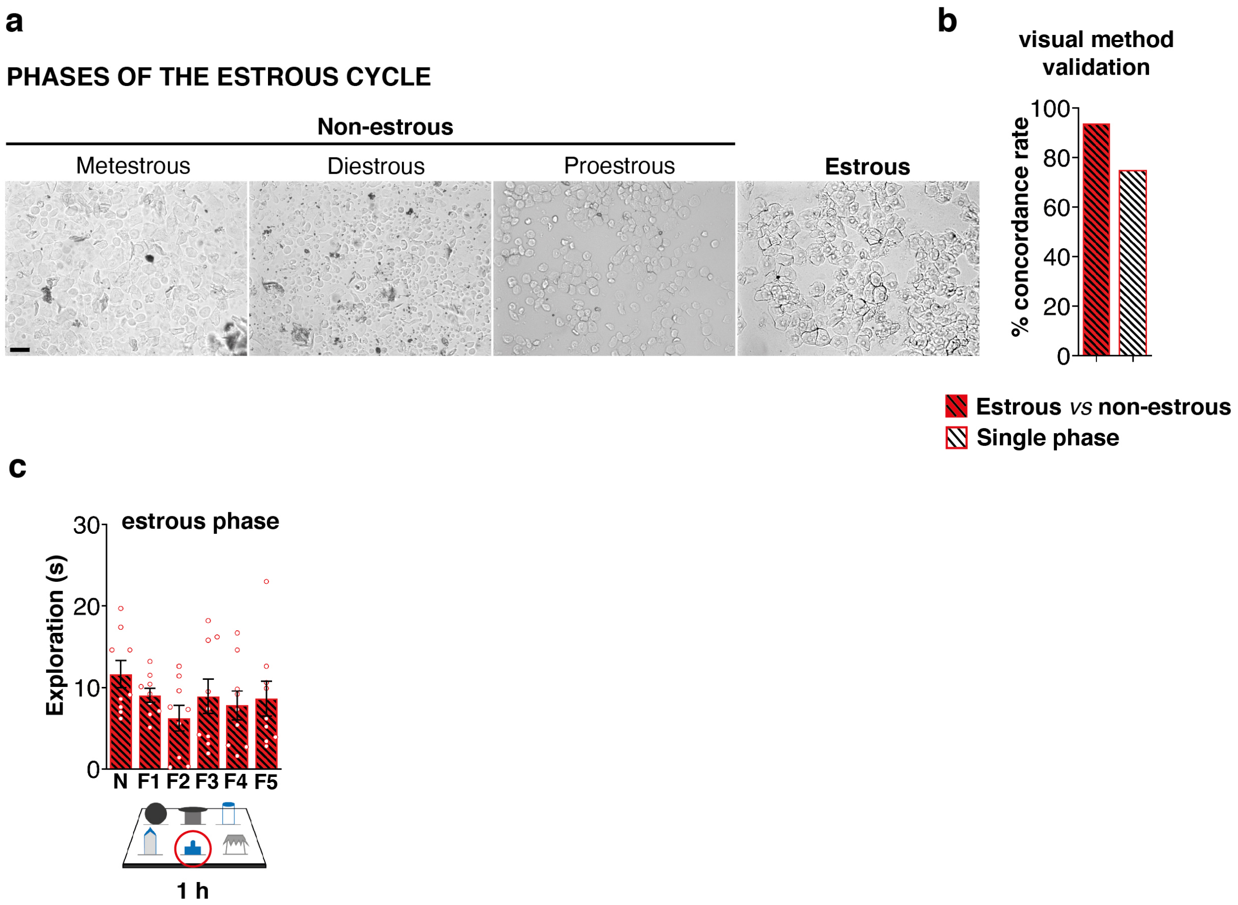


**Supplementary Figure 1.**

**(a)** Representative microphotographs of cellular type distribution from the different phases of the estrous cycle after vaginal smear collection. Scale bar: 50 µm.

**(b)** Bar charts indicate the percentage (%) concordance rate between visual inspection and vaginal smears analysis. The visual inspection proved more reliable in distinguishing estrous from non-estrous (n = 30/32), but not each estrous cycle (n = 24/32).

**(c)** Bar charts report exploration of the new object (circled) and each of the familiar ones (F). Females showed an impaired performance in the 6-DOT at 1 h delay under the estrous phase (n = 9) [one-way repeated measures ANOVA: $F_{5,40} = 1.203$; $p = 0.3253$]. Data are presented as mean values ± SEM.

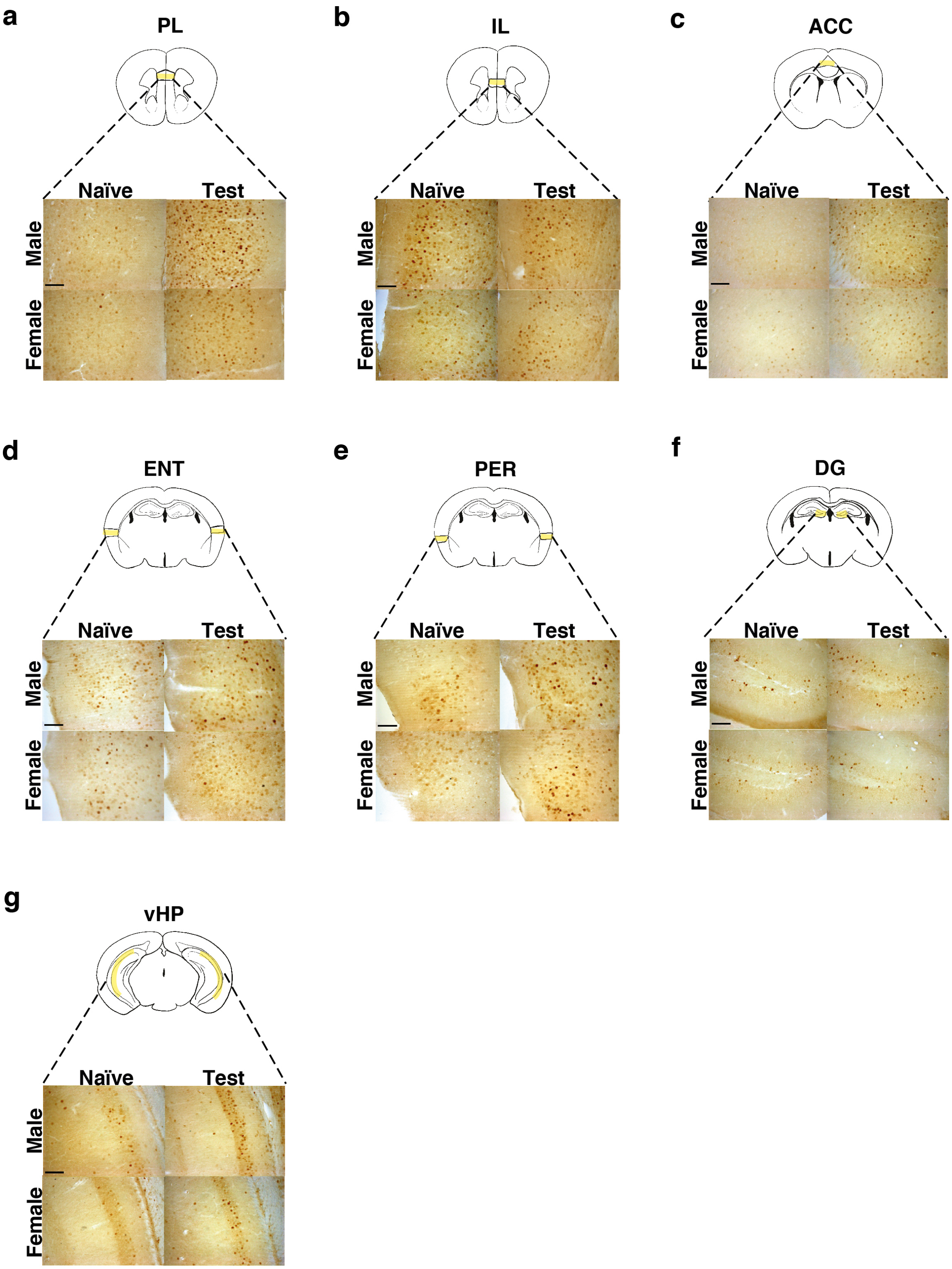
a
PL
Naïve
Test
Male
Female
b
IL
Naïve
Test
Male
Female
c
ACC
Naïve
Test
Male
Female
d
ENT
Naïve
Test
Male
Female
e
PER
Naïve
Test
Male
Female
f
DG
Naïve
Test
Male
Female
g
vHP
Naïve
Test
Male
Female

**Supplementary Figure 2.**

**(a-g).** Representative 20x magnification microphotographs of c-Fos immunohistochemical staining for PL, IL, ACC, ENT, PER, DG and vHP for each test condition and sex. Brown dots represent c-Fos$^+$ cells (Scale bar: 100 μm).

Statistical analyses for each brain region (bar charts in Fig. 2c):

The prelimbic cortex (PL) was found significantly activated only in male mice exposed to the 6-DOT [two-way ANOVA effect for: test $F_{1,21} = 5.008$, $p = 0.0362$, test *vs* sex $F_{1,21} = 5.318$, $p = 0.0314$].

No significant activation was found for the infralimbic cortex (IL) in respect to naïve or between sexes [two-way ANOVA effect for: test $F_{1,22} = 3.896$, $p = 0.0611$, test *vs* sex $F_{1,22} = 0.294$, $p = 0.5930$].

The anterior cingulate cortex (ACC) was found significantly activated only in male mice exposed to the 6-DOT [two-way ANOVA effect for: test $F_{1,27} = 28.160$, $p < 0.0001$, test *vs* sex $F_{1,27} = 8.080$, $p = 0.0084$].

The entorhinal cortex (ENT) was similarly activated in male and female mice in response to the 6-DOT [two-way ANOVA effect for: test $F_{1,19} = 15.280$, $p = 0.0009$, test *vs* sex $F_{1,19} = 1.005$, $p = 0.3287$; one-way ANOVA effect for test in males $F_{1,9} = 8.815$, $p = 0.0157$; one-way ANOVA effect for test in females $F_{1,10} = 9.252$, $p = 0.0124$].

The perirhinal cortex (PER) was similarly activated in male and female mice in response to the 6-DOT [two-way ANOVA effect for: test $F_{1,20} = 11.654$, $p = 0.0028$, test *vs* sex $F_{1,20} = 0.200$, $p = 0.6595$; one-way ANOVA effect for test in males $F_{1,9} = 6.474$, $p = 0.0315$; one-way ANOVA effect for test in females $F_{1,11} = 5.304$, $p = 0.0418$].

The dentate gyrus (DG) was found significantly activated only in male mice exposed to the 6-DOT [two-way ANOVA effect for: test $F_{1,27} = 32.037$, $p < 0.0001$, test *vs* sex $F_{1,27} = 9.873$, $p = 0.0040$].

The CA1-CA3 of the ventral hippocampus (vHP) was similarly activated in male and female mice in response to the 6-DOT [two-way ANOVA effect for: test $F_{1,15} = 25.588$, $p = 0.0001$, test *vs* sex $F_{1,15} = 2.658$, $p = 0.1239$; one-way ANOVA effect for test in males $F_{1,7} = 8.353$, $p = 0.0233$; one-way ANOVA effect for test in females $F_{1,8} = 18.773$, $p = 0.0025$].

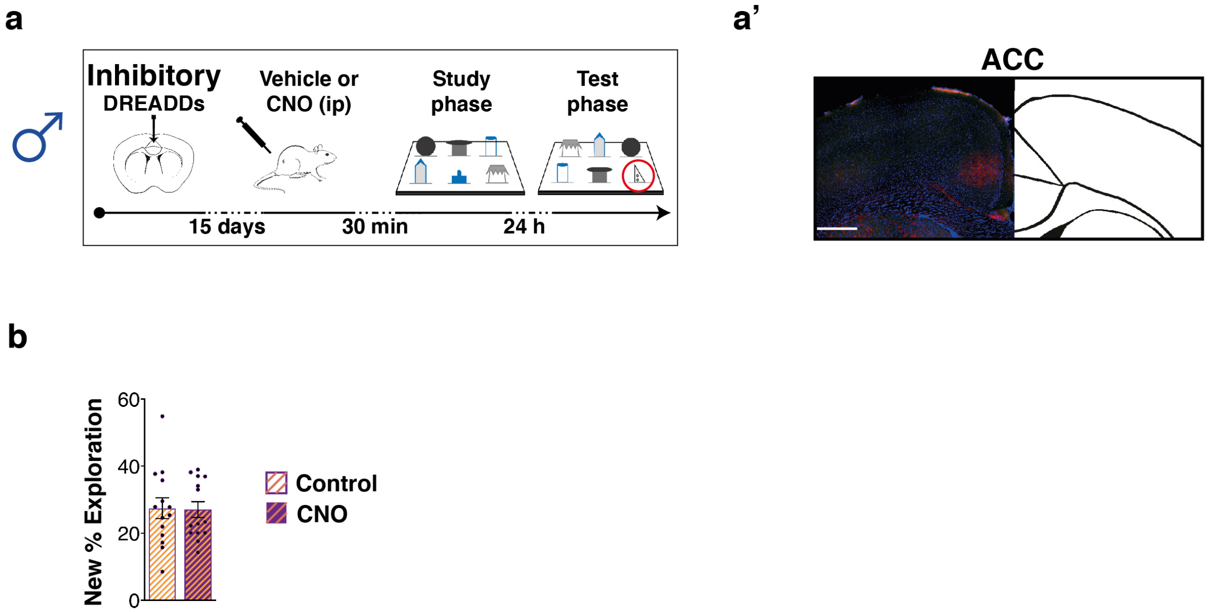


**Supplementary Figure 3.**

**(a)** Schematic of the experimental design showing the injection timeline of inhibitory DREADDs in the ACC of male mice. CNO or vehicle were injected 30 min before the task, to be active immediately after the end of the study phase, during early memory consolidation.

**(a')** Representative image of AAV expression in the ACC on the left and schematic of the ACC on the right; the microphotograph shows the inhibitory DREADDs signals through immunofluorescence analysis against hemagglutinin-tag (HA, red). Cells were counterstained with DAPI (blue) and the reporter mCitrine is shown in green. Scale bar 500 µm.

**(b)** No differences were found in the New object % exploration in CNO treated males ($n = 14$) compared to controls ($n = 14$) [Mann-Whitney test $p = 0.9820$]. Data are presented as mean values ± SEM.

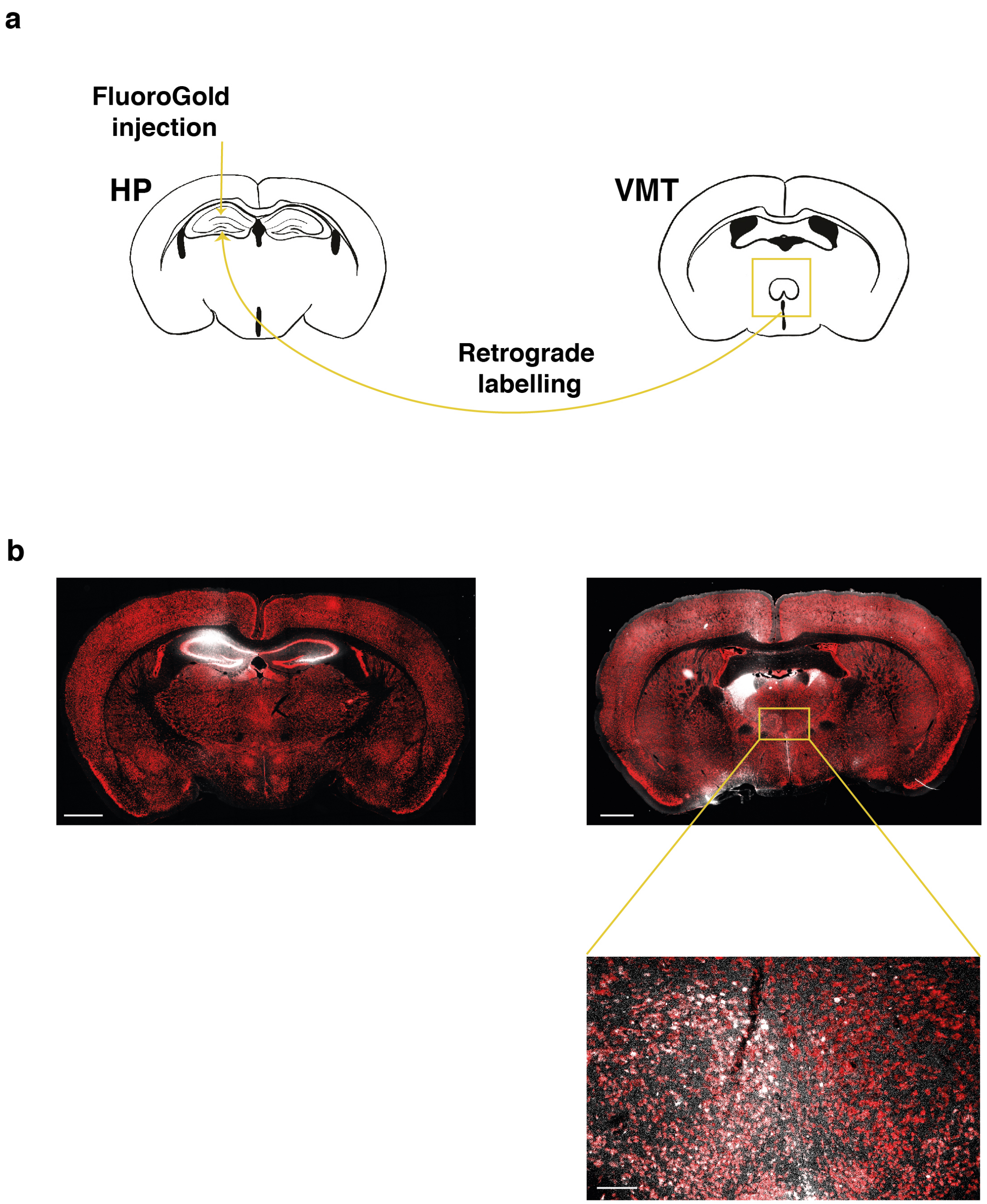


**Supplementary Figure 4**
**(a)** Schematic of the experimental design: the retrograde tracer FluoroGold was unilaterally injected in the HP of female mice and the retrograde labelling was qualitatively analyzed into the VMT.
**(b)** Representative fluorescence microscope images showing the unilateral injection site in the HP and FG labelled cells (white) in the ipsilateral VMT (scale bars in the mosaic slice reconstruction: 1000 µm; scale bar for VMT magnification: 100 µm).

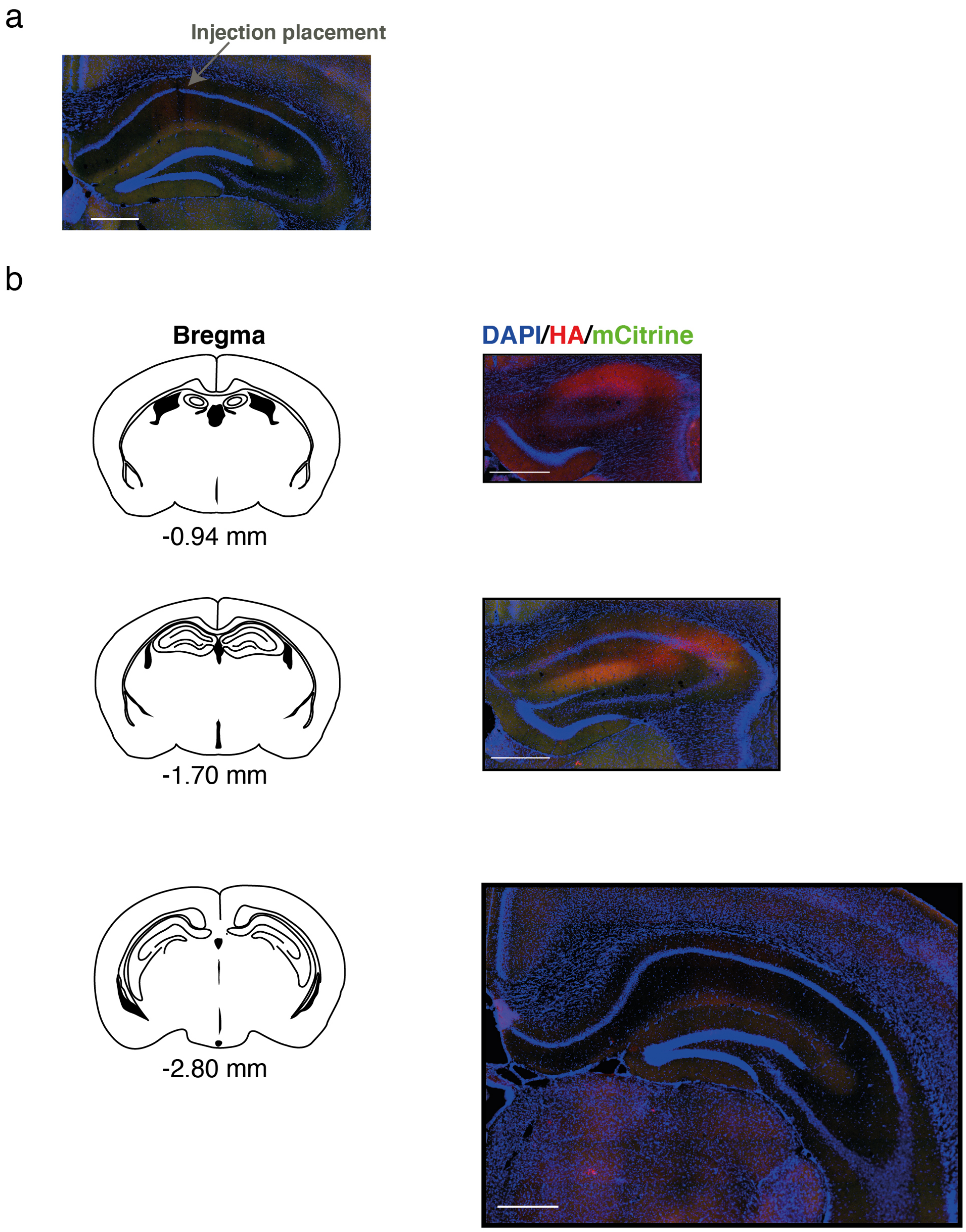


**Supplementary Figure 5**

**(a)** Microphotographs of the injection placement in the dHP for female mice injected with the activatory DREADDs (AAV 2/5-CaMKIIa-HA-rM3D(Gs)-IRES-mCitrine).

**(b)** On the left are reported the schematic representation of the antero-posterior coordinates from bregma of hippocampal slices where we identified the beginning (top) of the DREADDs expression, its major diffusion (center) and the end of the diffusion (bottom). On the right, microphotographs showing the adeno-associated virus signals through immunofluorescence analysis against hemagglutinin-tag (HA, red). Cells were counterstained with DAPI (blue) and the reporter mCitrine is shown in green. Scale bar 500 µm.

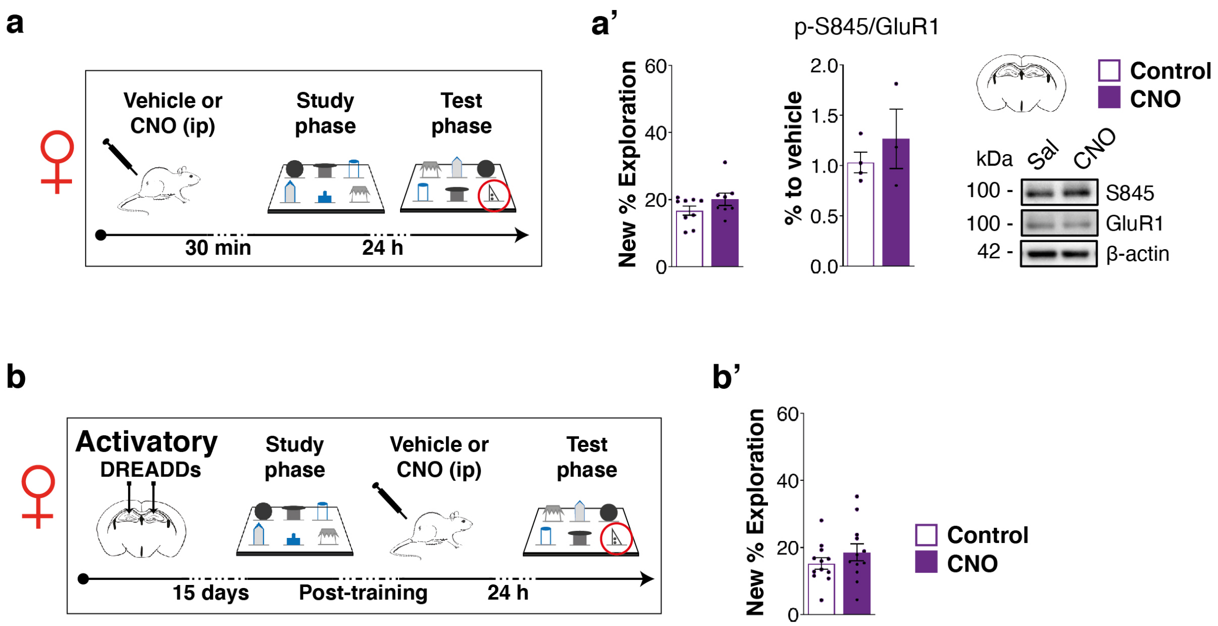


**Supplementary Figure 6**

**(a)** Schematic of the experimental design, where the red circled object represents the new item. A group of female mice received vehicle or CNO ip injection 30 min before the beginning of the 6-DOT and was tested 24 h later.

**(a')** Bar charts on the left represent the % New object exploration of control (n = 9) and CNO (n = 8) treated mice [Mann-Whitney test p = 0.1996]. Bar charts on the right represent western blot analysis of p-S845/GluR1 levels in the hippocampus of vehicle (4) and CNO (3) treated mice [one-way ANOVA for treatment $F_{1,5} = 0.731$, $p = 0.4315$]. No change was detected in GluR1 levels (saline: 100.0 ± 11.1; CNO: 73.4 ± 6.6; one-way ANOVA for treatment $F_{1,5} = 3.500$, $p = 0.1203$). Data are presented as mean values ± SEM.

**(b)** Schematic of the experimental design, where the red circled object represents the new item. Following activatory DREADDs injection in the dHP, mice received vehicle or CNO ip injection immediately after the study phase of the 6-DOT and were tested 24 h later.

**(b')** Bar charts represent the % New object exploration of vehicle (n = 12) and CNO (n = 12) treated mice [one-way ANOVA for treatment $F_{1,22} = 1.217$, $p = 0.2819$]. Data are presented as mean values ± SEM.

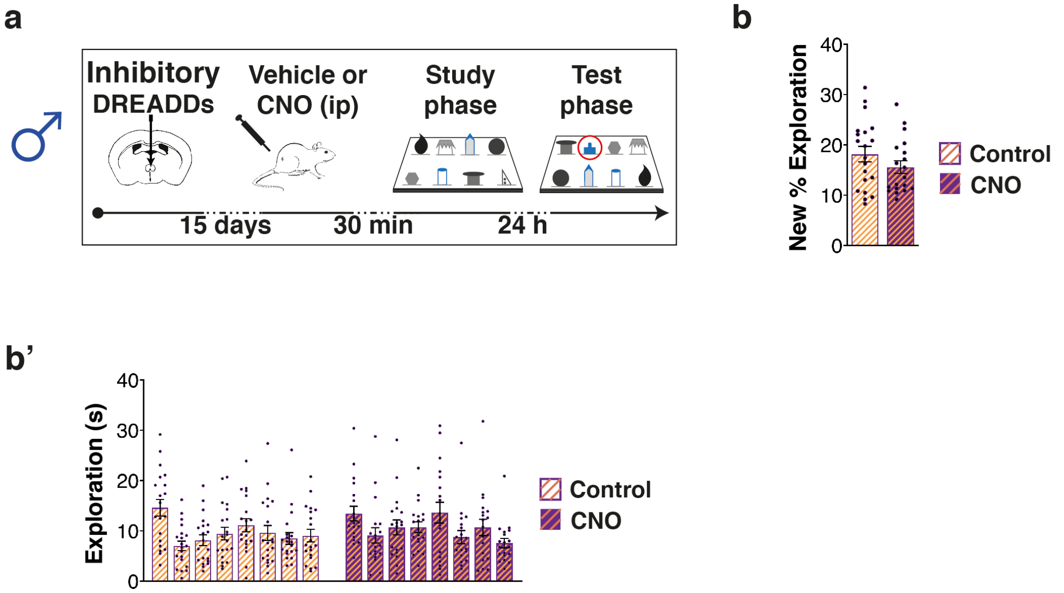


**Supplementary Figure 7**

**(a)** Schematic of the experimental design. Following the injection of inhibitory DREADDs (AAV-CaMKIIa-HA-rM4D(Gi)-IRES-mCitrine) in the VMT, male mice received vehicle or CNO ip injection 30 min before the beginning of the 8-DOT (overload condition) and were tested 24 h later.

**(b)** No differences were found in the new object % exploration in CNO treated males (n = 19) compared to controls (n = 20) [Mann-Whitney test p = 0.3363]. Data are presented as mean values ± SEM.

**(b')** The single object exploration for saline and CNO administered males injected with the inhibitory DREADDs in the VMT showed no differences between groups [two-way ANOVA for treatment and objects: effect for treatment $F_{1,37} = 1.155$; $p = 0.2895$, effect for objects $F_{7,259} = 4.769$; $p < 0.0001$, effect for treatment *vs* objects $F_{7,259} = 0.874$; $p = 0.5275$; one-way repeated measures ANOVA for objects: control: $F_{7,133} = 3.646$; $p = 0.0012$; CNO: $F_{7,126} = 2.2.2$; $p = 0.0383$]; neither group was able to discriminate the new object compared to all the familiar ones (Dunnett post-hoc test). Data are presented as mean values ± SEM.

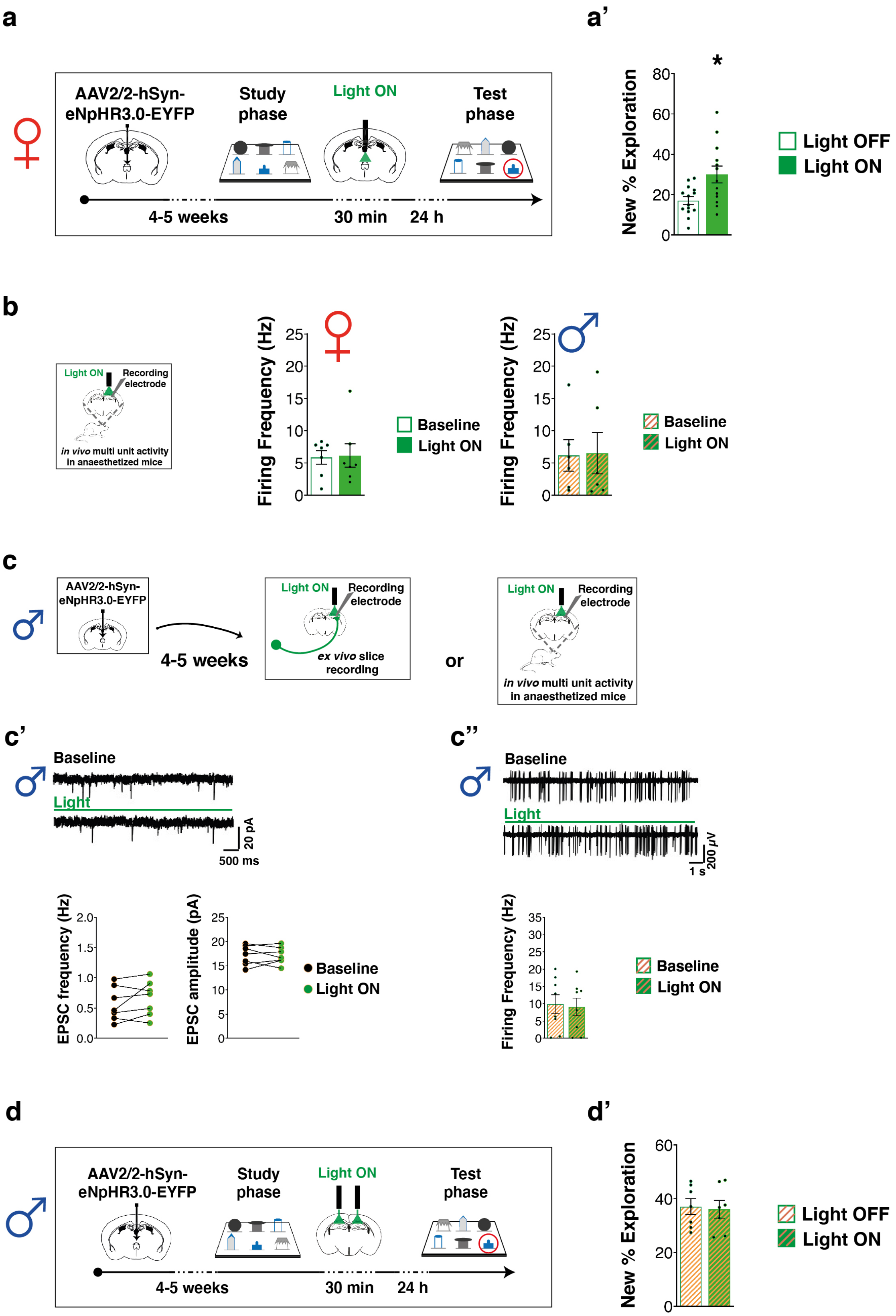
a
AAV2/2-hSyn-eNpHR3.0-EYFP
Study phase
Light ON
Test phase
4-5 weeks
30 min
24 h
a’
New % Exploration
*
Light OFF
Light ON
b
Light ON
Recording electrode
in vivo multi unit activity in anaesthetized mice
Firing Frequency (Hz)
Baseline
Light ON
c
AAV2/2-hSyn-eNpHR3.0-EYFP
4-5 weeks
Light ON
Recording electrode
ex vivo slice recording
or
in vivo multi unit activity in anaesthetized mice
c’
Baseline
Light
20 pA
500 ms
EPSC frequency (Hz)
EPSC amplitude (pA)
Baseline
Light ON
c’’
Baseline
Light
200 μV
1 s
Firing Frequency (Hz)
Baseline
Light ON
d
AAV2/2-hSyn-eNpHR3.0-EYFP
Study phase
Light ON
Test phase
4-5 weeks
30 min
24 h
d’
New % Exploration
Light OFF
Light ON

**Supplementary Figure 8**

**(a)** Schematic of the experimental design showing the timing of VMT post-training 30 min continuous inhibition; the new object in the phase is circled.

**(a')** Bar charts represent the % New object exploration in light off (n = 14) and light on (n = 13) female mice in the test conducted 24 h after training and VMT light stimulation [one-way ANOVA for light $F_{1,25} = 8.220$, $p = 0.0083$]. * $p < 0.05$ between groups. Data are presented as mean values ± SEM.

**(b)** Schematic of the *in vivo* hippocampal multiunit activity recordings of female and male naïve (no virus) mice under anesthesia. Bar charts represent spontaneous multiunit activity recorded in naïve female mice before (baseline: 5.9 ± 1.1 Hz) and during light stimulation (6.2 ± 1.8 Hz) [p= 0.9375, Wilcoxon test (n=7)] and naïve male mice before (baseline: 6.2 ± 2.5 Hz) and during light stimulation (6.5 ± 3.2 Hz) [p= 0.8438, Wilcoxon test (n=6)]. Data are presented as mean values ± SEM.

**(c)** Schematic of the experimental design for *ex-vivo* electrophysiological recordings in hippocampal slices and *in vivo* extracellular multiunit activity (MUA) in the pyramidal layer of the CA1 of anaesthetized male mice after AAV-hSyn-eNpHR3.0-EYFP injection in the VMT.

**(c')** On the top, representative traces from CA1 hippocampal neurons showing EPSCs in baseline and under photoinhibition (green bar). On the bottom, scatter plot showing EPSPs frequency and amplitude for males that showed no change after VMT-HP photoinhibition [n = 7 cells, 3 mice, EPSCs frequency: Wilcoxon test for light (Baseline, Light) p = 0.3750; EPSCs amplitude: Wilcoxon test for light (Baseline, Light) p = 0.9375].

**(c'')** On the top, representative traces of multiunit activity recorded from hippocampal CA1 pyramidal layer before and during photoinhibition (green bar). On the bottom, bar charts show no change in frequency activity in males between baseline (control) and under photoinhibition (light ON) [n = 8, Frequency: Wilcoxon test for light (Baseline, Light) p = 0.4609]. Data are presented as mean values ± SEM.

**(d)** Schematic of the experimental design showing the timing of VMT-dHP fibers post-training (30 min) continuous inhibition in male mice.

**(d')** Bar charts represent the New object % exploration in light OFF (n = 17) and light ON (n = 13) males in the test conducted 24 h after post-training VMT-dHP photoinhibition. No effect was found on the performance at test phase in male mice [One-way ANOVA for light (Light OFF, Light ON): $F_{1,28} = 0.684$; p = 0.4152]. Data are presented as mean values ± SEM.

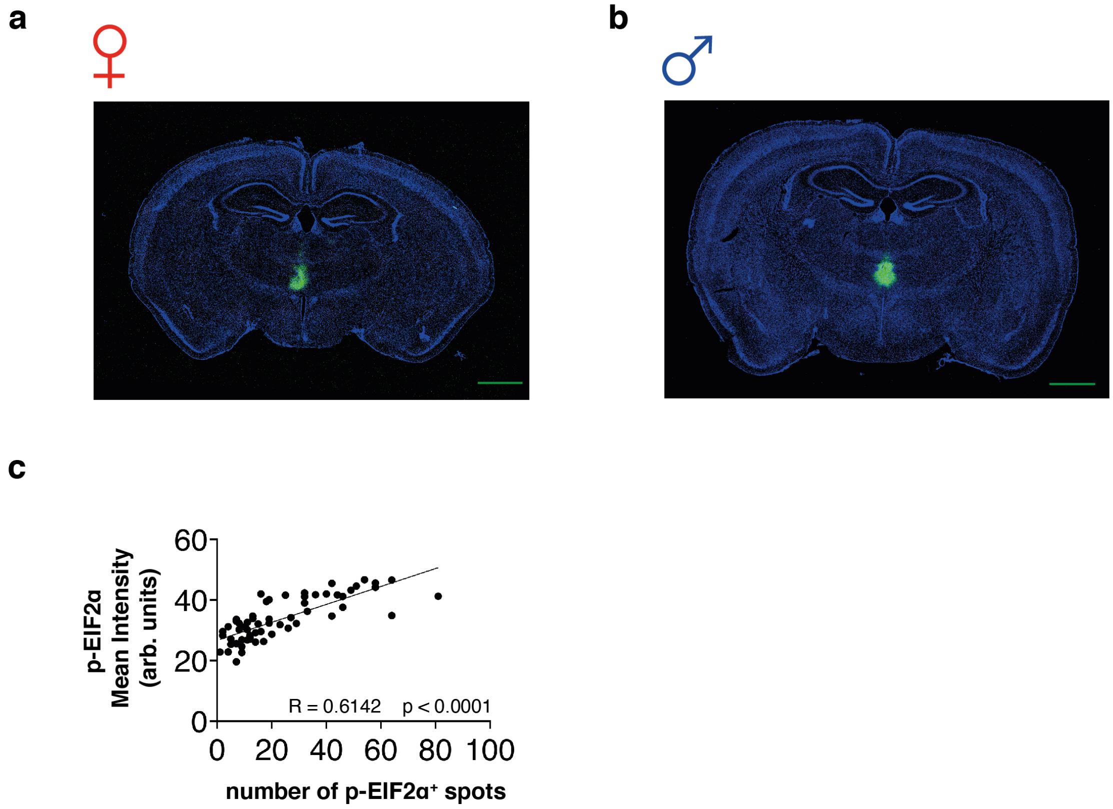


**Supplementary Figure 9**

**(a-b)** Representative images of the AAV injection in the VMT of a female (a) and a male (b) mouse from the VMT-HP optogenetic photoinhibition experiment; scale bars: 1000 µm.

**(c)** Correlation coefficient between the number of p-EIF2α$^+$ spots and p-EIF2α mean intensity measured in excitatory neurons (GAD67$^-$) per ROIs.

**Supplementary Table 1**

The table reports the mean total objects exploration ± SEM during the study phase of the Different/Identical Object Task (DOT/IOT) for different load conditions (3, 4, 6 or 9 objects) and the number of subjects *per* group.

| Number of Objects | Task | Delay | Total exploration study phase (sec ± SEM) | N |
|---|---|---|---|---|
| 3 | IOT | 1 min | 33.6 ± 1.8 | 7 |
| | DOT | 1 min | 104.5 ± 3.8 | 15 |
| | DOT | 24 h | 78.0 ± 12.2 | 7 |
| 4 | IOT | 1 min | 35.8 ± 1.2 | 8 |
| | DOT | 1 min | 128.9 ± 9.3 | 7 |
| | DOT | 24 h | 35.7 ± 2.5 | 14 |
| 6 | IOT | 1 min | 35.4 ± 1.4 | 18 |
| | DOT | 1 min | 195.1 ± 7.2 | 18 |
| | DOT | 24 h | 145.1 ± 11.8 | 14 |
| 9 | IOT | 1 min | 31.1 ± 1.3 | 9 |
| | DOT | 1 min | 267.3 ± 14.7 | 12 |

**Supplementary Table 2**

The table reports the mean total objects exploration ± SEM during the study phase of the 6-DOT/IOT with 1 min and 24 h retention interval and the number of subjects *per* group. In the high memory load condition (6-DOT), females show a higher level of total objects exploration compared to males. Moreover, females spend significantly less time than males to reach the cut-off exploration of 210 sec in the 6-DOT. * $p < 0.05$ between sexes (one-way ANOVA).

| Retention interval | Task | Sex | Total exploration study phase (sec ± SEM) | Time to reach the cut-off exploration in the study phase (min ± SEM) | N |
|---|---|---|---|---|---|
| 1 min and 24 h | 6-IOT | male | 33.8 ± 0.6 | 2.1 ± 0.2 | 23 |
| | | female | 35.9 ± 1.2 | 2.0 ± 0.1 | 23 |
| | 6-DOT | male | 154.8 ± 6.9 | 9.8 ± 0.1 | 33 |
| | | female | 180.4 ± 7.4* | 9.0 ± 0.2* | 34 |

**Supplementary Table 3**

c-Fos basal level in male and female naïve mice used for normalization of data in Fig. 2C. Data are expressed as percentage of cells ± SEM per groups of slices on the anteroposterior extension of each brain structure analyzed. No significant difference was found between groups (one-way ANOVA).

| Brain structure | Sex | % cells | n | p value<br>F value |
|---|---|---|---|---|
| Prelimbic cortex | Male | 83.1 ± 35.5 | 5 | p = 0.7551 |
| | Female | 96.1 ± 19.1 | 5 | $F_{1,8} = 0.104$ |
| Infralimbic cortex | Male | 97.3 ± 49.9 | 5 | p = 0.9798 |
| | Female | 98.9 ± 37.3 | 6 | $F_{1,9} = 0.001$ |
| Anterior cingulate cortex | Male | 100.9 ± 35.4 | 6 | p = 0.8825 |
| | Female | 93.2 ± 36.2 | 7 | $F_{1,11} = 0.023$ |
| Ventral midline thalamus | Male | 91.03 ± 14.5 | 9 | p = 0.5364 |
| | Female | 76.7 ± 16.4 | 12 | $F_{1,19} = 0.396$ |
| Entorhinal cortex | Male | 90.2 ± 45.3 | 3 | p = 0.9059 |
| | Female | 100.0 ± 47.6 | 7 | $F_{1,8} = 0.015$ |
| Perirhinal cortex | Male | 91.7 ± 46.4 | 3 | p = 0.6364 |
| | Female | 100.0 ± 45.7 | 7 | $F_{1,8} = 0.012$ |
| Dorsal hippocampus | Male | 100.0 ± 93.7 | 5 | p = 1.0 |
| | Female | 100.0 ± 56.8 | 8 | $F_{1,11} = 0.000$ |
| Dentate gyrus | Male | 100.0 ± 64.1 | 5 | p = 1.0 |
| | Female | 100.0 ± 31.3 | 8 | $F_{1,11} = 0.000$ |
| Ventral hippocampus | Male | 83.3 ± 74.5 | 3 | p = 0.8140 |
| | Female | 100.0 ± 32.6 | 6 | $F_{1,7} = 0.060$ |

**Supplementary Table 4**

The table reports the mean total distance ± SEM in the familiarization trial, the mean total objects exploration ± SEM during the study and test phase, and the mean total exploration ± SEM of familiar objects during the test phase of the 6-DOT with 24 h retention interval. * $p < 0.05$ between groups (one-way ANOVA).

| Figure | Treatment | Total distance T1 (m ± SEM) | Total exploration T2 (sec ± SEM) | Total exploration T3 (sec ± SEM) | Total familiar objects exploration T3 (sec ± SEM) |
|---|---|---|---|---|---|
| 3b | Control | 30.2 ± 4.6 | 146.1 ± 15.2 | 45.0 ± 5.2 | 35.8 ± 3.8 |
| | CNO | 28.3 ± 2.8 | 136.2 ± 9.6 | 39.8 ± 3.7 | 28.1 ± 3.8 |
| 3c | Control | 48.4 ± 4.9 | 128.9 ± 12.8 | 44.1 ± 4.6 | 35.8 ± 4.2 |
| | CNO | 42.7 ± 3.3 | 123.2 ± 10.9 | 46.2 ± 3.6 | 32.4 ± 2.1 |
| 3d’ | Control | 33.4 ± 4.2 | 84.7 ± 7.1 | 35.3 ± 3.7 | 26.1 ± 2.8 |
| | CNO | 38.0 ± 3.8 | 73.3 ± 9.7 | 32.8 ± 2.0 | 27.0 ± 1.9 |
| 4c’ | Light OFF | 41.0 ± 4.1 | 92.2 ± 12.2 | 33.6 ± 2.5 | 27.4 ± 2.2* |
| | Light ON | 33.0 ± 5.2 | 92.5 ± 10.1 | 28.0 ± 2.9 | 18.9 ± 2.2 |
| S3b | Control | 46.5 ± 3.5 | 58.2 ± 5.9 | 37.4 ± 4.0 | 26.3 ± 2.5 |
| | CNO | 46.5 ± 3.4 | 52.0 ± 4.2 | 39.3 ± 3.0 | 28.6 ± 2.5 |
| S7b | Control | 36.7 ± 3.3 | 69.3 ± 9.1 | 77.5 ± 4.7 | 62.9 ± 3.6 |
| | CNO | 36.2 ± 2.7 | 79.5 ± 7.7 | 84.7 ± 4.6 | 71.2 ± 3.9 |
| S8a’ | Light OFF | 35.3 ± 3.2 | 87.6 ± 6.6 | 36.4 ± 2.0 | 30.3 ± 2.0 |
| | Light ON | 30.1 ± 3.1 | 82.9 ± 8.4 | 34.5 ± 2.6 | 24.7 ± 2.3 |
| S8d’ | Light OFF | 38.2 ± 3.7 | 80.0 ± 7.0 | 26.6 ± 2.4 | 17.5 ± 1.6 |
| | Light ON | 36.2 ± 2.5 | 85.7 ± 8.5 | 29.9 ± 2.1 | 20.8 ± 1.6 |

**Supplementary Table 5**

The statistical analyses of the figures in the main text are reported here. The number of excluded animals because of procedural testing problems, outliers, wrong injection placement or no viral expression is reported in round brackets.

| Figure | Statistical test | Sample size |
|---|---|---|
| **1a** | **Object exploration**: Repeated measures ANOVA for 3-IOT at 1min: $F_{2,12} = 9.926$; $p = 0.0029$ | Females = 7 |
| | **Object exploration**: Repeated measures ANOVA for 3-DOT at 1 min: $F_{2,28} = 23.876$; $p < 0.0001$ | Females = 15 |
| **1b** | **Object exploration**: Repeated measures ANOVA for 4-IOT at 1min: $F_{3,21} = 15.541$; $p < 0.0001$ | Females = 8 |
| | **Object exploration**: Repeated measures ANOVA for 4-DOT at 1 min: $F_{3,18} = 12.140$; $p = 0.0001$ | Females = 7 |
| **1c** | **Object exploration**: Repeated measures ANOVA for 6-IOT at 1min: $F_{5,85} = 20.019$; $p < 0.0001$ | Females = 18 |
| | **Object exploration**: Repeated measures ANOVA for 6-DOT at 1 min: $F_{5,85} =12.163$; $p < 0.0001$ | Females = 18 |
| **1d** | **Object exploration**: Repeated measures ANOVA for 9-IOT at 1min: $F_{8,64} = 8.450$; $p < 0.0001$ | Females = 9 |
| | **Object exploration**: Repeated measures ANOVA for 9-DOT at 1 min: $F_{8,88} = 1.193$; $p = 0.3124$ | Females = 12 |
| **1e** | **Object exploration**: Repeated measures ANOVA for 3-DOT at 24h: $F_{2,12} = 13.364$; $p = 0.0009$ | Females = 7 (1) |
| **1f** | **Object exploration**: Repeated measures ANOVA for 4-DOT at 24h: $F_{3,39} = 13.347$; $p < 0.0001$ | Females = 14 |
| **1g** | **Object exploration**: Repeated measures ANOVA for 6-DOT at 24h: $F_{5,65} = 0.271$; $p < 0.9273$ | Females = 14 (2) |
| **1h** | **Object exploration**: Repeated measures ANOVA for 6-DOT at 1h: $F_{5,45} = 1.585$; $p = 0.1838$ | Females = 10 |
| **1i'** | **% New**: Three-way ANOVA for sex (male, female) *vs* treatment (control, interference) *vs* load (6-IOT, 6-DOT): effect for sex $F_{1,48} = 5.228$; $p = 0.0267$; effect for load $F_{1,48} = 15.792$; $p = 0.0002$.<br>**% New (Males-6-IOT):** One-way ANOVA for treatment (control, interference): $F_{1,13} = 0.403$; $p = 0.5366$.<br>**% New (Females-6-IOT):** One-way ANOVA for treatment (control, interference): $F_{1,11} = 0.021$; $p = 0.8871$.<br>**% New (Males-6-DOT):** One-way ANOVA for treatment (control, interference): $F_{1,12} = 5.541$; $p = 0.0365$.<br>**% New (Females-6-DOT):** One-way ANOVA for treatment (control, interference): $F_{1,12} = 0.028$; $p = 0.8704$. | Males 6-IOT:<br>Control = 7<br>Interference = 8<br>Males 6-DOT:<br>Control = 7 (1)<br>Interference = 7 (1)<br>Females 6-IOT:<br>Control = 6 (2)<br>Interference = 7 (1)<br>Females 6-DOT:<br>Control = 7<br>Interference = 7 (2) |

| | | |
|---|---|---|
| **2a** | **% New**: Three-way ANOVA for sex (male, female) *vs* delay (1 min, 24 h) *vs* load (6-IOT, 6-DOT): effect for sex *vs* delay $F_{1,105} = 8.954$; $p = 0.0035$.<br>Two-way ANOVA for sex (male, female) *vs* load (6-IOT, 6-DOT) at 1 min delay: effect for sex $F_{1,54} = 0.498$; $p = 0.4833$; effect for sex *vs* load $F_{1,54} = 0.233$; $p = 0.6389$.<br>Two-way ANOVA for sex (male, female) *vs* load (6-IOT, 6-DOT) at 24 h delay: effect for sex $F_{1,51} = 14.124$; $p = 0.0004$; effect for load $F_{1,51} = 9.501$; $p = 0.0033$; effect for sex *vs* load $F_{1,51} = 1.124$; $p = 0.2940$.<br>One-way ANOVA for load (6-IOT, 6-DOT) at 24h in female mice: $F_{1,26} = 8.045$; $p = 0.0087$.<br>One-way ANOVA for sex (male, female) at 24 h in the 6-DOT: $F_{1,37} = 29.043$; $p < 0.0001$ | Males 6-IOT:<br>1 min = 15<br>24 h = 8<br>Males 6-DOT:<br>1 min = 14<br>24 h = 19<br>Females 6-IOT:<br>1 min = 15<br>24 h = 8<br>Females 6-DOT:<br>1 min = 14<br>24 h = 20 |
| **2c** | **% c-Fos cells to naïve in dHP:** Two-way ANOVA for sex (male, female) *vs* test (naïve, 6-DOT): effect for load $F_{1,23} = 10.678$; $p = 0.0034$: effect for load *vs* sex $F_{1,23} = 4.328$; $p = 0.0488$ | Males:<br>Naïve = 5<br>6-DOT = 7 (2)<br>Females:<br>Naïve = 8<br>6-DOT = 7 (1) |
| | **% c-Fos cells to naïve in VMT:** Two-way ANOVA for sex (male, female) *vs* test (naïve, 6-DOT): effect for load $F_{1,37} = 29.227$; $p < 0.0001$; effect for load *vs* sex $F_{1,37} = 9.625$; $p = 0.0037$ | Males:<br>Naïve = 9<br>6-DOT = 11 (2)<br>Females:<br>Naïve = 12<br>6-DOT = 9 |
| **2d** | **% c-Fos cells to naïve in dHP**: Two-way ANOVA for sex (male, female) *vs* load (naïve, 6-IOT, 3-DOT, 6-DOT): effect for sex $F_{1,54} = 5.885$; $p = 0.0186$; effect for load $F_{3,54} = 16.976$; $p < 0.0001$; effect for sex *vs* load $F_{3,54} = 6.456$; $p = 0.0008$ | Males:<br>Naïve = 11<br>6-IOT = 5<br>3-DOT = 6<br>6-DOT = 6 (2)<br>Females:<br>Naïve = 14<br>6-IOT = 5<br>3-DOT = 6<br>6-DOT = 9 (1) |
| **2e** | **% c-Fos cells to naïve in VMT**: Two-way ANOVA for sex (male, female) *vs* load (naïve, 6-IOT, 3-DOT, 6-DOT): effect for sex $F_{1,55} = 11.128$; $p = 0.\ 0015$; effect for load $F_{3,55} = 15.164$; $p < 0.0001$; effect for sex *vs* load $F_{3,55} = 4.613$; $p = 0.0060$ | Males:<br>Naïve = 9<br>6-IOT = 5<br>3-DOT = 6<br>6-DOT = 11 (2)<br>Females:<br>Naïve = 12<br>6-IOT = 5<br>3-DOT = 6<br>6-DOT = 9 |
| **3b** | **% New**: One-way ANOVA for treatment (Control, CNO): $F_{1,22} = 5.563$; $p = 0.0276$ | Females:<br>Control = 12 (2)<br>CNO = 12 (2) |

| | | |
|---|---|---|
| **3b'** | **% p-S845/GluR1**: One-way ANOVA for treatment (Control, CNO): Mann-Whitney test = 0.0480 | Females:<br>Control = 5<br>CNO = 7 |
| | **% GluR1** [Control: 100.0 ± 9.8; CNO: 114.4 ± 12.2]: One-way ANOVA for treatment (Control, CNO): $F_{1,10} = 0.737$; p = 0.4106 | |
| **3c** | **% New**: One-way ANOVA for treatment (Control, CNO): $F_{1,22} = 9.112$; p = 0.0063 | Females:<br>Control = 12 (3)<br>CNO = 12 (4) |
| **3c'** | **% p-S845/GluR1**: One-way ANOVA for treatment (Control, CNO): treatment $F_{1,9} = 5.341$; p = 0.0462 | Females:<br>Control = 6<br>CNO = 5 |
| | **% GluR1** [Control: 100.0 ± 10.8; CNO: 307.2 ± 95.9]: One-way ANOVA for treatment (Control, CNO) $F_{1,9} = 5.628$; p = 0.0417 | |
| **3d'** | **% New**: One-way ANOVA for treatment (Control, CNO): $F_{1,24} = 5.406$; p = 0.0289 | Males:<br>Control = 12 (2)<br>CNO = 14 (1) |
| | **% p-S845**: One-way ANOVA for treatment (Control, CNO): Mann-Whitney test = 0.0357 | Males:<br>Control = 3<br>CNO = 5 |
| | **% GluR1**: One-way ANOVA for treatment (Control, CNO): Mann-Whitney test = 0.0357 | Males:<br>Control = 3<br>CNO = 5 |
| | **% p-S845/GluR1** [Control: 100.0 ± 22.8; CNO: 62.1 ± 4.7]: One-way ANOVA for treatment (Control, CNO): $F_{1,6} = 4.538$; p = 0.0772 | Males:<br>Control = 3<br>CNO = 5 |
| **4b'** | **EPSCs frequency**: Wilcoxon test for light (Baseline, Light) p = 0.0039<br>**EPSCs amplitude**: Wilcoxon test for light (Baseline, Light) p = 0.0391 | Females = 3<br>Cells = 9 |
| **4b''** | **Frequency**: Wilcoxon test for light (Baseline, Light) p= 0.0195 | Females = 10 |
| **4c'** | **% New**: One-way ANOVA for light (Light OFF, Light ON): $F_{1,27} = 10.114$; p = 0.0037 | Females:<br>light off = 14 (6)<br>light on = 15 (2) |
| **4d'** | **c-Fos$^{+}$ cells on Light OFF in py**: One-way ANOVA for light (Light OFF, Light ON): $F_{1,17} = 7.961$; p = 0.0118 | Females slices:<br>light off = 9<br>light on = 10 |
| **4f'** | **c-Fos$^{+}$ cells on Light OFF in so**: One-way ANOVA for light (Light OFF, Light ON): $F_{1,12} = 7.260$; p = 0.0195.<br>**c-Fos$^{+}$/GAD$^{+}$ cells on Light OFF in so**: One-way ANOVA for light (Light OFF, Light ON): $F_{1,12} = 1.359$; p = 0.2664.<br>**% c-Fos$^{+}$/GAD$^{+}$ cells on total c-fos$^{+}$ cells in so**: One-way ANOVA for light (Light OFF, Light ON): $F_{1,12} = 6.064$; p = 0.0299. | Females slices:<br>light off = 6<br>light on = 8 |
| **4g'** | **p-EIF2α/EIF2α**: One-way ANOVA for light (Light OFF, Light ON): Mann-Whitney test = 0.0049 | Females slices:<br>light off = 9<br>light on = 8 |
